# Deciphering auditory neuroprogenitor stemness for large-scale neuronal screening

**DOI:** 10.64898/2026.10.09.757611

**Authors:** Francis Rousset, Lorenzo Pepe, Moein Sarhadi, Sylvain Lemeille, Stéphanie Sgroi, Lucie Oberhauser, Dimitrios Daskalou, Sten Ilmjärv, Simon Braun, Pascal Senn

**Author notes:** equal contribution (Lorenzo Pepe, Moein Sarhadi).

## Abstract

Sensorineural hearing loss can arise from damage to sensory hair cells, auditory neurons or their synaptic connections, with auditory neuropathy and cochlear synaptopathy highlighting the importance of neural integrity for hearing. The mammalian cochlea has limited capacity to regenerate lost auditory neurons. Understanding the molecular programs that sustain cochlear progenitor expansion and neurogenic competence could support neuronal regeneration and provide renewable models for auditory drug discovery. We previously established that WNT activation combined with dual SMAD inhibition (CHIR-DS) supports scalable expansion and neuronal differentiation of mouse auditory neuroprogenitors (ANPGs). Here, we characterize the transcriptional, chromatin and metabolic changes associated with the induced state. RNA-seq and ATAC-seq identified reduced Id1 and Id2 expression and increased accessibility at proneural basic helix-loop-helix motifs, alongside transcriptional enrichment of translation, mTOR and glucose-metabolism programs. Compared with growth-factor controls, CHIR-DS-treated ANPGs showed a slight increase in glycolysis at early passage. However, after long-term expansion, both glycolysis and mitochondrial respiration were markedly enhanced. Human fetal ANPGs also responded to CHIR-DS with increased sphere growth and retained neural differentiation capacity, providing proof of concept beyond the mouse system. Finally, neurons derived from stemness-induced ANPGs (siANs) supported a 384-well reporter assay and a 2,560-compound pilot screen that nominated candidate neurite-outgrowth enhancers. Together, these findings validate the pathway basis of ANPG stemness induction, characterize the resulting molecular and metabolic state, and demonstrate human feasibility and a neuronal screening application.

**HIGHLIGHTS:**

- CHIR-DS reduces Id1/Id2 expression and increases proneural bHLH motif accessibility
- Long-term ANPG cultures show increased glycolytic and respiratory activity
- Human fetal ANPGs expand under CHIR-DS and retain neural differentiation capacity
- ANPG-derived neurons support a 384-well pilot screen of 2,560 compounds

## INTRODUCTION

Hearing depends on the integrity of both mechanosensory hair cells and their auditory neural connections. Type I spiral ganglion neurons (SGNs) receive glutamatergic input from inner hair cells at ribbon synapses and convey temporally structured action-potential trains to the cochlear nucleus [1]. Their tonotopic organization preserves frequency information, while differences in firing thresholds, spontaneous activity and dynamic range contribute to encoding sound intensity. Single-cell studies have resolved molecularly distinct type I SGN subtypes with characteristic ion-channel profiles and peripheral connectivity [2–4]. The integrity of this neural pathway can be compromised independently of hair-cell survival: in mice, noise exposure causes persistent loss of afferent terminals and delayed neuronal degeneration despite recovery of hearing thresholds [5]. Lost SGNs are not effectively replaced in the mature mammalian cochlea, and restoring hair cells alone cannot rebuild an absent neural pathway [6]. Cochlear implants likewise depend on surviving auditory nerve fibres to convey electrical stimulation [7]. These considerations make neuronal preservation, synaptic repair and neuronal replacement complementary objectives for hearing restoration.

Preclinical studies support both cell replacement and recruitment of resident cochlear cells as approaches to neural repair. Human embryonic stem-cell-derived otic neuroprogenitors engrafted, differentiated and improved auditory-evoked response thresholds after transplantation into a gerbil model of auditory neuropathy [8]. Cochlear glia provide a candidate cellular source for neuronal replacement. SOX2-positive auditory nerve glia display neural stem/progenitor properties in culture, and cochlear SOX2-positive glia can generate neurons under defined reprogramming conditions [6, 9]. In neonatal mice, Neurog1 and Neurod1 expression converts cochlear glia into SGN-like neurons [10]. WNT/β-catenin activation also enhances neuronal differentiation of lineage-traced FZD10-positive cochlear glia following SGN injury in early postnatal mice [11]. These findings motivate the identification of molecular regulators that sustain cochlear neurogenic competence.

Auditory neuroprogenitors (ANPGs) isolated from the postnatal spiral ganglion provide a tissue-derived culture system for investigating such regulators. These neurosphere-forming populations retain neuronal and glial differentiation potential, but their expansion under conventional growth-factor conditions is limited to a few passages [12–15]. We previously identified Phoenix ANPGs from A/J mice, which exhibit sustained intrinsic self-renewal and neuronal differentiation capacity[14]. Guided by their molecular profile, we subsequently established scalable expansion of C57BL/6 ANPGs through WNT activation with CHIR99021 combined with dual SMAD inhibition using LDN193189 and SB431542 (CHIR-DS). This published protocol supports long-term ANPG amplification while preserving the capacity to generate functional neurons [15]. While scalable ANPG production is already established, the molecular changes through which CHIR-DS induces this expandable neurogenic state remain incompletely defined.

Neural progenitor expansion and neuronal differentiation competence depend on coordinated regulation of cell-cycle activity and neurogenic programs. In other neural progenitor systems, proneural basic helix-loop-helix factors can support both proliferation and neuronal commitment, with their effects depending on expression dynamics [16, 17]. Epigenomic profiling has identified distinct enhancer programs controlling neural stem-cell quiescence [18, 19], while single-cell transcriptomics has linked progenitor activation to changes in translational capacity, metabolism and energy use [20]. These observations provide a rationale for examining the transcriptional, chromatin and metabolic responses of ANPGs to CHIR-DS.

Understanding the molecular response to CHIR-DS could identify candidate entry points for stimulating endogenous auditory neurogenesis and inform progenitor-based replacement strategies for auditory neuropathy. The established expansion platform also provides a renewable source of neurons for investigating auditory neuronal injury and screening compounds that promote neuronal survival or neurite outgrowth[14, 15]. Extending these conditions to human progenitors is an important step toward renewable human in vitro models, building on previous demonstrations that human fetal cochlear progenitors can be expanded and differentiated into neurons [21].

Here, we characterize the CHIR-DS-induced state in mouse ANPGs using RNA-seq and ATAC-seq during early induction and extracellular flux measurements during early and long-term culture, complemented by multiplex genetic silencing of the targeted signaling pathways. We also assess CHIR-DS responsiveness and neural differentiation in human fetal ANPGs. Finally, we use neurons derived from stemness-induced ANPGs (siANs) in a 384-well pilot screen of 2,560 compounds. Together, these analyses seek to identify molecular programs relevant to auditory neuronal regeneration and extend the use of renewable ANPG-derived neuronal models.

## RESULTS

### ANPG stemness induction is accompanied by transcriptional and chromatin remodeling

To investigate the molecular changes associated with ANPG stemness induction, we compared four culture conditions selected for their different effects on progenitor expansion (**Figure 1A, B**). Growth factors alone (GF) provided the reference condition, while CHIR99021 (CHIR) was used to examine the response to WNT activation. CHIR was combined with either the BMP receptor inhibitor K02288 (CHIR+BMP) or LDN193189 and SB431542 to achieve dual inhibition of BMP and TGFβ–SMAD signaling (CHIR-DS). These combined conditions are labelled +WNT-BMP and +WNT-DS, respectively, in **Figure 1**. CHIR+BMP and CHIR-DS promote greater ANPG expansion than GF or CHIR alone, and CHIR-DS supports sustained propagation while preserving neuronal differentiation capacity, as previously established [15]. This comparison allowed us to distinguish responses to WNT activation alone from molecular changes shared by the two conditions that enhance ANPG expansion.

**Figure 1.**
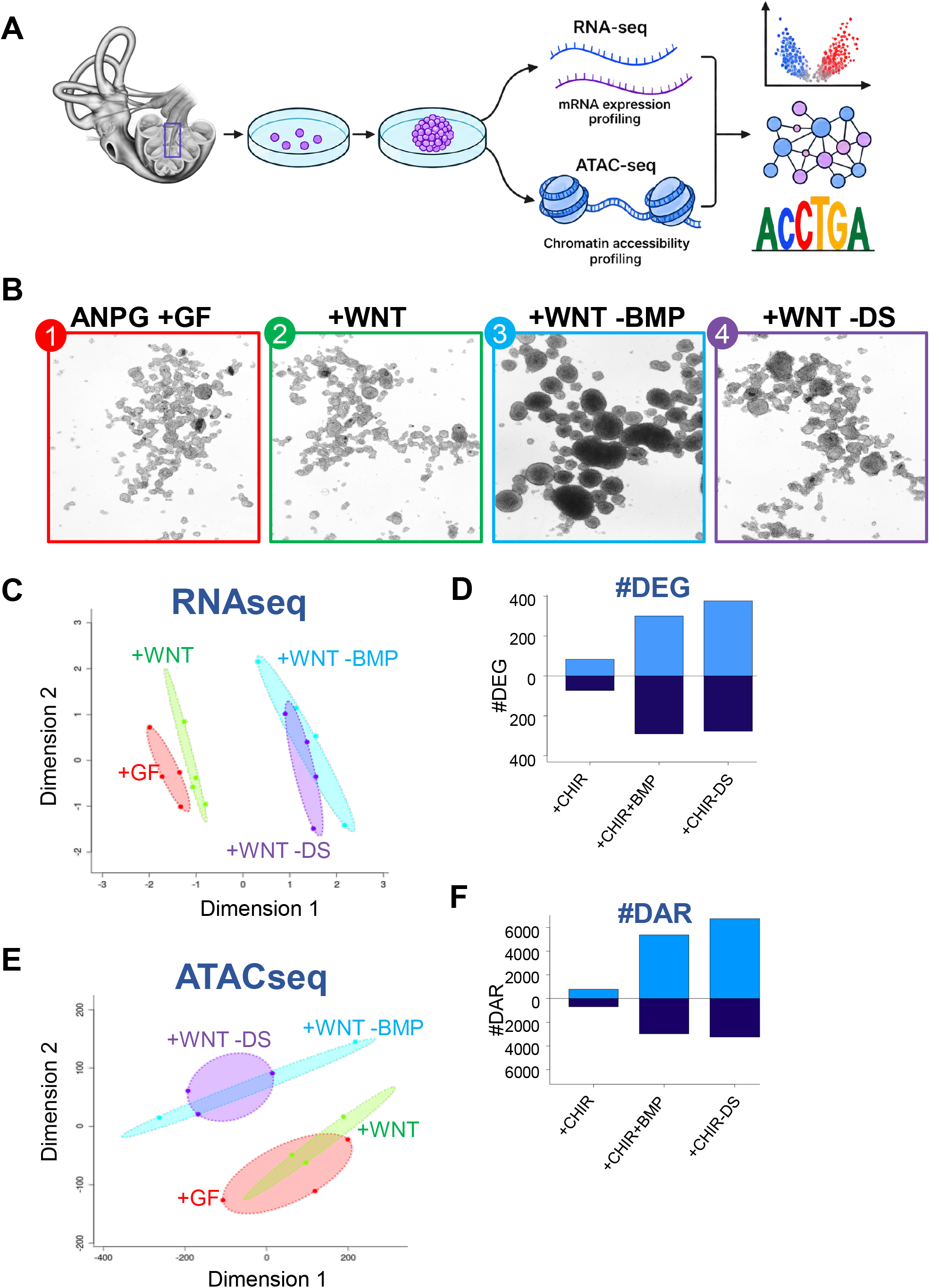
Early transcriptional and chromatin accessibility changes during ANPG stemness induction. **(A)** Schematic of the experimental design. Auditory neuroprogenitors (ANPGs) were isolated from the postnatal mouse cochlea, cultured as primary neurospheres, and exposed for 10 days to control conditions or stemness-inducing conditions. RNA-seq and ATAC-seq were then performed to define early transcriptional and chromatin accessibility changes associated with reprogramming. **(B)** Representative bright-field images of primary ANPG neurospheres cultured under the indicated conditions: control, +WNT, +WNT-BMP, and +WNT-DS. **(C)** RNA-seq dimensionality reduction plot showing segregation of control and +WNT samples from +WNT-BMP and +WNT-DS samples. **(D)** Number of differentially expressed genes (DEGs) identified in +WNT, +WNT-BMP, and +WNT-DS conditions relative to control. **(E)** ATAC-seq dimensionality reduction plot showing chromatin accessibility differences across conditions, with +WNT-BMP and +WNT-DS clustering apart from control and +WNT. **(F)** Number of differentially accessible regions (DARs) identified in +WNT, +WNT-BMP, and +WNT-DS conditions relative to control. ANPGs were isolated from postnatal day 5 (P5) C57BL/6 mice and cultured for 10 days under the following conditions: control (growth factors only), +WNT (CHIR99021), +WNT-BMP (CHIR99021 + K02288), and +WNT-DS (CHIR99021 + SB431542 + LDN193189). Data represent 3 -4 biological replicates.

ANPGs isolated from postnatal day 5 C57BL/6J mice were cultured under these conditions for 10 days before parallel RNA-seq and ATAC-seq analyses to assess early changes in gene expression and chromatin accessibility, respectively (**Figure 1A**). Multidimensional scaling separated the GF and CHIR groups from the CHIR+BMP and CHIR-DS groups in both datasets (**Figure 1C, E**). Relative to GF, the combined treatments produced more differentially expressed genes and differentially accessible chromatin regions than CHIR alone (**Figure 1D, F**).

We also used genetic silencing to assess the contribution of the signaling nodes targeted by CHIR-DS. Synthetic miRNA minigenes based on our multiplex silencing technology [22] were designed to target *Alk2*, *Alk5* and *Gsk3b*, corresponding to BMP, TGFβ and GSK3β-dependent regulation of WNT signaling, respectively (**Supplementary Figure 1A**). Combined targeting increased neurosphere area after 30 days to the range observed with CHIR-DS, and genetically targeted cultures maintained sphere formation during long-term expansion (**Supplementary Figure 1B, C**). This response provides genetic support for the contribution of the targeted signaling nodes to ANPG expansion.

### Stemness induction is associated with metabolic remodeling

Gene set enrichment analysis revealed coordinated changes in signaling and metabolic programs across the treatment comparisons. GF cultures were enriched for calcium, cAMP and GPCR signaling, extracellular matrix interactions and electron-transport pathways (**Figure 2A**). Treatment-associated enrichment included translation, glucose metabolism, stress responses and mTORC1-associated gene sets (**Figure 2B**). In the direct CHIR-DS versus GF comparison, enrichment analyses identified WNT signaling and pluripotency, HIF1 signaling, mTOR signaling and translation among the treatment-associated programs (**Figure 2C-F**). Differential expression analysis identified 277 genes more highly expressed in GF and 376 in CHIR-DS. Genes upregulated under CHIR-DS included Crabp1, Wnt7b and Pdgfra, whereas Id1, Id2, Id3, Hes1 and Dlk1 were downregulated (**Figure 2G**).

**Figure 2.**
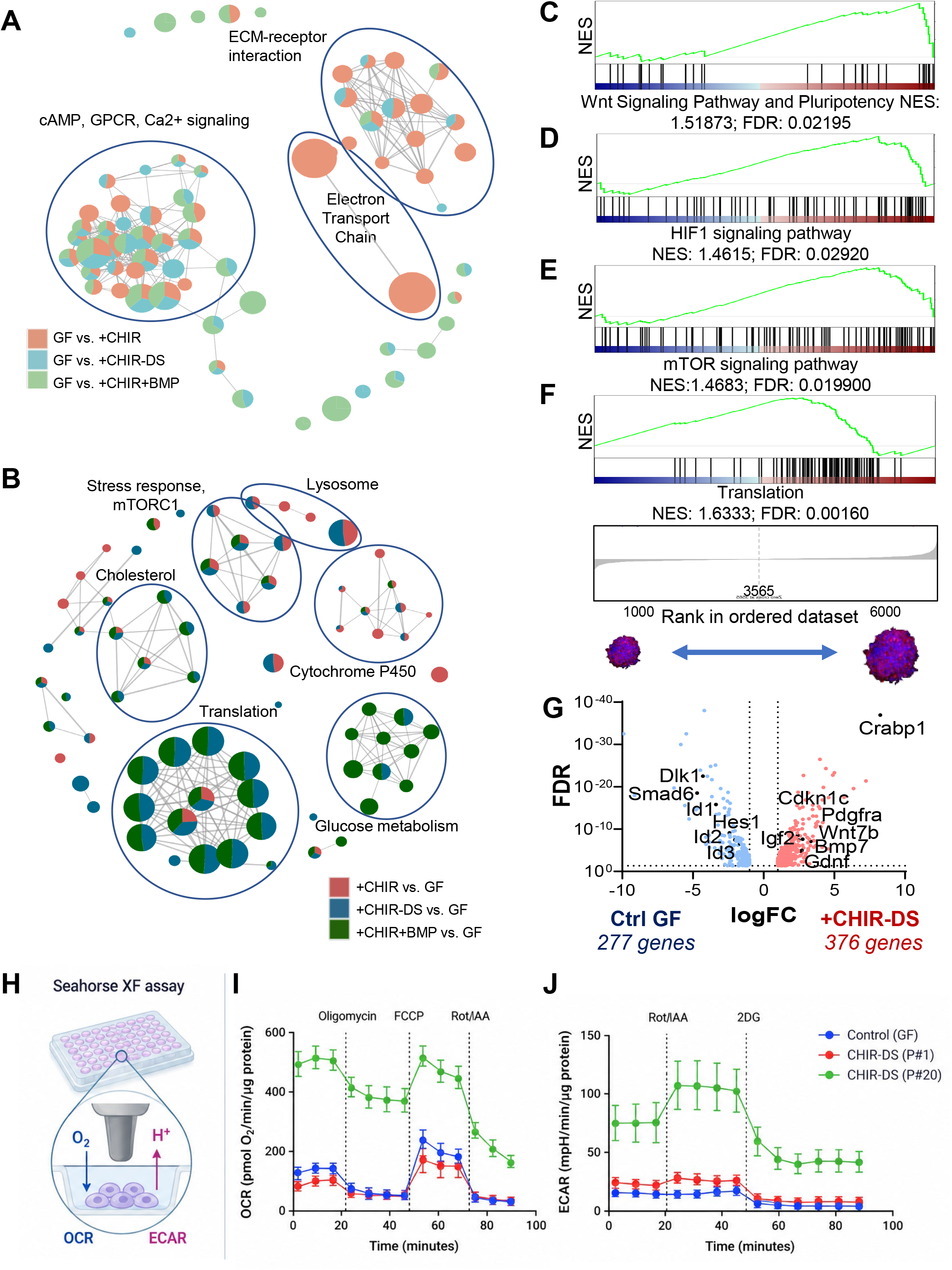
Metabolic shift of ANPGs upon stemness induction. **(A-B)** Gene Set Enrichment Analysis (GSEA) network maps summarizing pathway-level changes across stemness inducing conditions. **(A)** Pathways enriched in GF relative to +CHIR, +CHIR-BMP, and +CHIR-DS. **(B)** Pathways enriched in +CHIR, +CHIR-BMP, and +CHIR-DS relative to GF. Node size reflects gene set size, and edges indicate shared genes between pathways. **(C-F)** Representative GSEA enrichment plots from the +CHIR-DS versus GF comparison, showing enrichment of Wnt signaling/pluripotency (**C**; NES = 1.51873, FDR = 0.02195), HIF1 signaling (**D**; NES = 1.4615, FDR = 0.02920), mTOR signaling (**E**; NES = 1.4683, FDR = 0.01990), and translation (**F**; NES = 1.6333, FDR = 0.00160). The lower bars indicate gene rank positions within the ordered dataset. **(G)** Volcano plot of differentially expressed genes comparing low-stemness control ANPGs (C57BL/6J; 277 genes) and high-stemness +CHIR-DS ANPGs (376 genes). Selected genes associated with stemness induction, WNT signaling, and growth-related programs are highlighted. **(H)** Schematic of the Seahorse XF assay used to assess metabolic activity in ANPGs, measuring oxygen consumption rate (OCR) and extracellular acidification rate (ECAR). **(I)** OCR profiles of GF control, early-passage +CHIR-DS (P#1), and long-term +CHIR-DS (P#20) ANPGs during Seahorse analysis following sequential injection of oligomycin, FCCP, and rotenone/antimycin A (Rot/AA). **(J)** ECAR profiles of the same conditions during Seahorse analysis following sequential injection of rotenone/antimycin A and 2-deoxyglucose (2DG).Seahorse data are normalized to protein content and expressed as pmol O2/min/µg protein (OCR) or mpH/min/µg protein (ECAR). Together, these data indicate that stemness induction is accompanied by broad transcriptional reprogramming and a metabolic shift consistent with enhanced glycolytic and respiratory activity.

To determine whether these transcriptional changes were accompanied by altered metabolic activity, we performed Seahorse extracellular flux analysis in GF control ANPGs, newly induced CHIR-DS cultures after 10 days (passage 1), and long-term CHIR-DS cultures at passage 20. Oxygen consumption rate (OCR) and extracellular acidification rate (ECAR) were normalized to the BCA protein measurement for each well (**Figure 2H-J**).

In the mitochondrial stress test, oligomycin reduced OCR in all groups, identifying an ATP-linked component of respiration (**Figure 2I**). FCCP increased OCR above basal levels in GF and passage 1 cultures, revealing respiratory reserve. Passage 20 cultures displayed markedly higher basal OCR, but FCCP-stimulated OCR remained close to basal levels, indicating limited additional respiratory reserve under these assay conditions. Rotenone and antimycin A markedly reduced OCR, confirming a substantial mitochondrial contribution to oxygen consumption.

Basal ECAR was modestly increased at passage 1 and markedly elevated at passage 20 compared with GF controls (**Figure 2J**). Rotenone and antimycin A further increased ECAR in CHIR-DS cultures, most prominently at passage 20, consistent with compensatory glycolysis following mitochondrial inhibition. Subsequent addition of 2-deoxyglucose strongly reduced ECAR, confirming a substantial glycolytic contribution to extracellular acidification.

Together, these profiles reveal a modest early glycolytic response followed by a pronounced elevation of both glycolytic activity and mitochondrial respiration during long-term expansion. The combination of high basal respiratory activity and the ability to further recruit glycolysis during mitochondrial inhibition is consistent with sustained metabolic demand and an adaptive response to metabolic challenge in expanding ANPG cultures.

### Reduced Id and Hes1 expression accompanies increased accessibility at proneural bHLH motifs

The downregulation of *Id* genes and *Hes1* prompted us to examine chromatin changes at loci associated with repression of proneural programs. Accessibility at *Id1* and *Id2* decreased under CHIR+BMP and CHIR-DS, in agreement with their reduced transcript abundance (**Figure 3C, D**). Reduced accessibility was also observed at *Hes1* and the corepressor locus *Tle1* (**Figure 3F, G**). These changes involve two complementary components of proneural regulation: ID-mediated inhibition of E-protein partners and HES1-associated transcriptional repression (**Figure 3E, H**).

**Figure 3.**
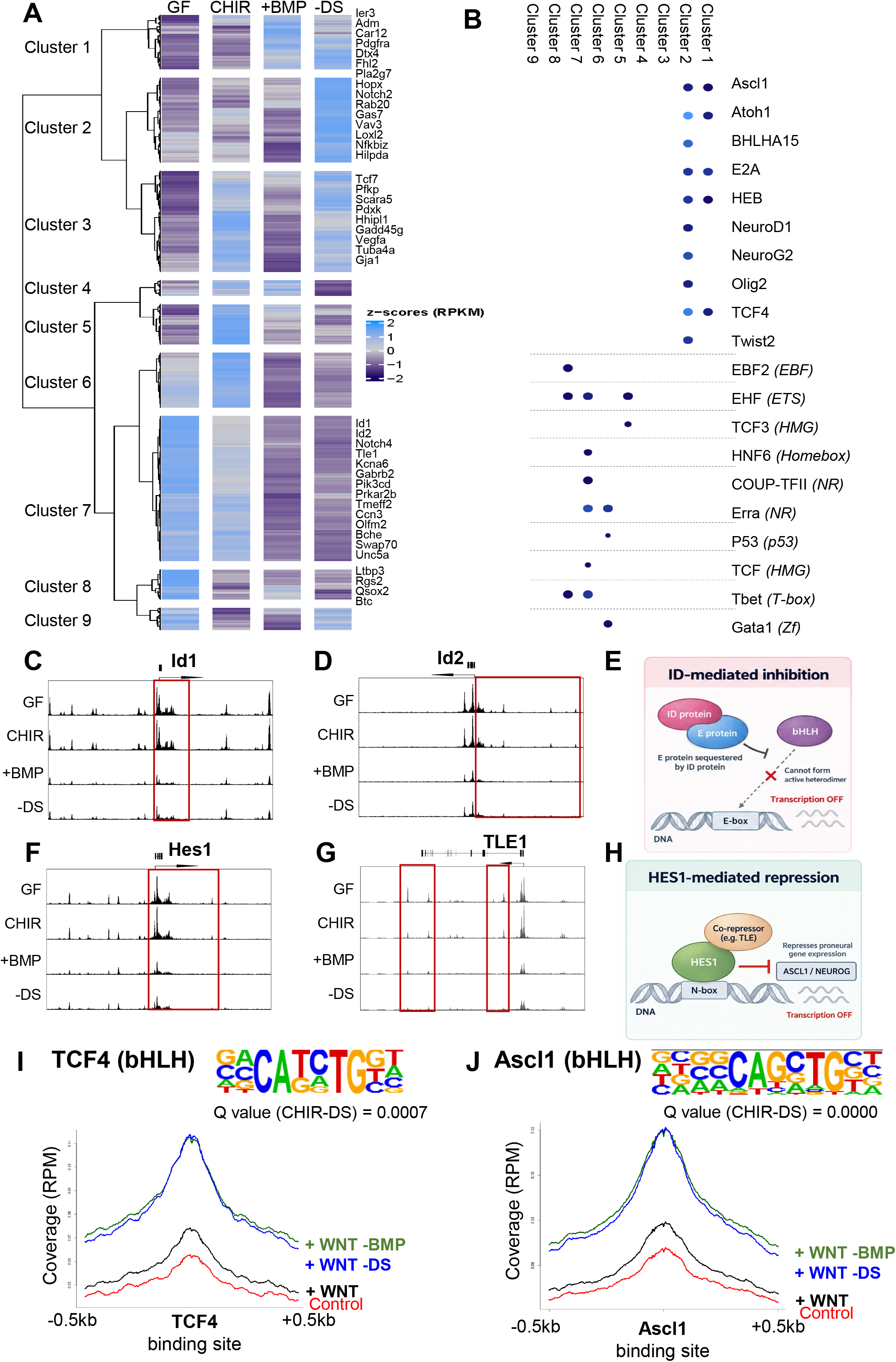
Chromatin accessibility remodeling of proneural bHLH motifs during stemness induction. **(A)** Heatmap of differentially accessible regions identified by ATAC-seq across GF, CHIR, +BMP, and -DS conditions. Nine clusters with distinct accessibility dynamics are shown. Representative genes associated with each cluster are indicated on the right. **(B)** Transcription factor motif enrichment across accessibility clusters. Clusters associated with stemness-inducing conditions are enriched for proneural bHLH motifs, including *Ascl1, Atoh1, NeuroD1, NeuroG2,* and *TCF4*, whereas clusters associated with low-stemness conditions show enrichment for alternative motifs, including EBF, ETS, nuclear receptor, HMG, and other lineage-or state-associated factors. **(C-D)** Representative ATAC-seq coverage tracks at the *Id1* (**C**) and *Id2* (**D**) loci across GF, CHIR, +BMP, and -DS conditions. Highlighted regions denote regulatory elements showing reduced accessibility under stemness-inducing conditions, consistent with repression of *Id*-associated inhibitory programs. **(E)** Schematic of ID-mediated inhibition. ID proteins antagonize bHLH transcription factor activity by sequestering E proteins, thereby preventing formation of active bHLH/E-protein complexes and suppressing transcription of target genes. **(F-G)** Representative ATAC-seq coverage tracks at the *Hes1* (**F**) and *TLE1* (**G**) loci across conditions. Highlighted regions indicate differentially accessible regulatory elements, consistent with remodeling of the Notch-HES-TLE repressive network during stemness induction. **(H)** Schematic of HES1-mediated repression. HES1 represses proneural gene expression through recruitment of co-repressors such as TLE, thereby maintaining transcriptional inhibition of neuronal differentiation programs. **(I-J)** Motif logos and aggregate ATAC-seq signal centered on genomic occurrences of the TCF4/E2-2 **(I)** and Ascl1 **(J)** motifs. Accessibility around both motifs is increased under stemness-inducing conditions relative to GF and CHIR, consistent with increased accessibility at regulatory sequences containing proneural bHLH motifs. Q values for motif enrichment are indicated.

Hierarchical clustering of differentially accessible regions identified nine clusters with distinct responses to treatment (**Figure 3A**). Regions in clusters 1 and 2 were preferentially accessible under stemness-inducing conditions, whereas clusters 7 and 8 showed greater accessibility in GF cultures. Motif analysis revealed enrichment of proneural bHLH motifs in the induced-state clusters, including ASCL1, ATOH1, NEUROD1 and NEUROG2, together with the E-protein motifs E2A, HEB and TCF4/E2-2 (Figure 3B). Aggregate ATAC-seq signal around genomic TCF4/E2-2 and ASCL1 motif occurrences was lowest in GF cultures, intermediate with CHIR alone and highest with CHIR+BMP or CHIR-DS (**Figure 3I, J**). These results associate *Id/Hes1* downregulation with increased chromatin accessibility at proneural bHLH motifs during ANPG stemness induction.

Given the changes at *Hes1*, we also assessed the effect of pharmacological Notch inhibition on ANPG expansion. The γ-secretase inhibitors avagacestat and DAPT did not significantly enhance sphere growth beyond the corresponding CHIR or CHIR-DS conditions (**Supplementary Figure 2A-C**). Cultures retained Nestin and SOX2 expression and generated βIII-tubulin-positive and S100-positive cells after differentiation (**Supplementary Figure 2D-H** and **D**′**-H**′). Additional Notch inhibition therefore provided no detectable expansion advantage under the conditions tested.

### CHIR-DS supports sustained expansion and neural differentiation of human fetal ANPGs

We next examined whether the protocol established in mouse ANPGs could be applied to human fetal cultures. Over 32 days, CHIR increased sphere growth relative to GF, and the largest response was observed with CHIR-DS. Dual SMAD inhibition alone did not significantly increase growth over GF (**Figure 4A-C**). At passage 5, CHIR-DS cultures continued to form abundant spheres, whereas GF cultures showed limited propagation (**Figure 4D-F**).

**Figure 4.**
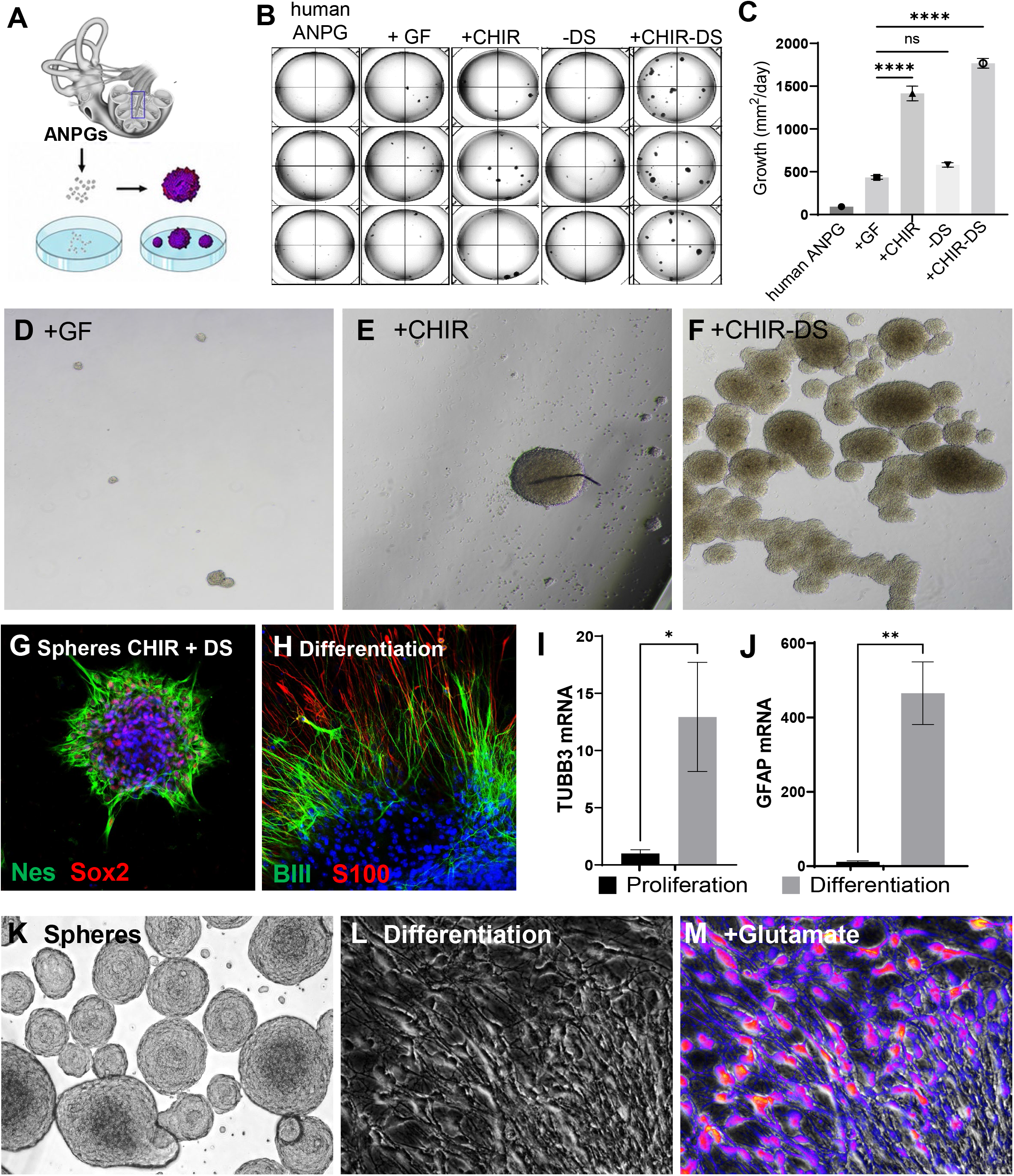
Stemness induction enables sustained expansion and neural differentiation of human fetal auditory neuroprogenitors. **(A)** Schematic of the human auditory neuroprogenitor (ANPG) sphere-growth assay. Human fetal ANPGs were isolated from the cochlea, seeded at 10,000 cells per well in 96-well plates, and cultured under control or stemness-inducing conditions. **(B)** Representative macroscopic images of ANPG cultures after 32 days under basal conditions, growth factors (+GF), WNT activation (+CHIR), dual-SMAD inhibition (-DS), or combined WNT activation and dual-SMAD inhibition (+CHIR-DS). CHIR99021 was used at 3 µM; dual-SMAD inhibition consisted of LDN193189 (0.5 µM) and SB431542 (10 µM). **(C)** Quantification of ANPG growth over 32 days, expressed as mean sphere growth per day. Combined WNT activation and dual-SMAD inhibition markedly increased ANPG expansion. Data are presented as mean ± SEM; statistical significance is indicated in the graph. **(D-F)** Representative bright-field images of human fetal ANPGs at passage 5 cultured with growth factors alone (**D**), +CHIR (**E**), or +CHIR-DS (**F**). +CHIR-DS supported robust and sustained sphere formation, whereas cultures maintained with growth factors alone showed limited propagation. **(G-H)** Phenotypic characterization of +CHIR-DS-expanded human ANPGs. **(G)** Immunofluorescence of an expanded neurosphere showing expression of the neural progenitor markers Nestin (Nes, green) and SOX2 (red), with nuclei counterstained with DAPI (blue). **(H)** Following 7 days of differentiation after withdrawal of growth and reprogramming factors, cultures expressed the neuronal marker βIII-tubulin (BIII, green) and the glial marker S100 (red). **(I-J)** Expression of neuronal and glial differentiation markers before and after differentiation. Differentiation significantly increased TUBB3 (**I**) and GFAP (**J**) mRNA levels compared with proliferating ANPG cultures. Data are presented as mean ± SEM; statistical significance is indicated in the graphs. **(K-M)** Neural differentiation and functional assessment of human fetal ANPGs. **(K)** Representative bright-field image of human fetal ANPG spheres generated from a 12-week post-conception cochlea. **(L)** Representative morphology after 7 days of differentiation on a Matrigel-coated surface following withdrawal of growth factors. **(M)** Representative image of differentiated ANPG-derived cells loaded with the Ca²⁺-sensitive indicator Fluo-8 and stimulated with 100 µM glutamate. The pseudocolored signal indicates glutamate-evoked intracellular Ca²⁺ mobilization, supporting acquisition of excitatory neuronal responsiveness.

Expanded cultures expressed the progenitor markers Nestin and SOX2 (**Figure 4G**). Following 7 days of differentiation after withdrawal of growth and reprogramming factors, cultures contained βIII-tubulin-positive neuronal cells and S100-positive cells, accompanied by increased *TUBB3* and *GFAP* transcript levels (**Figure 4H-J**). The transition from spheres to adherent cultures with neuronal processes was also apparent by transmitted-light microscopy (**Figure 4K, L**). Differentiated cells loaded with Fluo-8 exhibited intracellular calcium mobilization following stimulation with 100 μM glutamate (**Figure 4M**). These findings provide proof of concept that CHIR-DS supports sustained human fetal ANPG expansion while preserving neural differentiation capacity and the ability to generate glutamate-responsive cells.

### High-throughput calcium imaging compares glutamate responses across differentiation conditions

To identify differentiation conditions suitable for functional neuronal assays, we compared glutamate responsiveness in neurons derived from stemness-induced mouse ANPGs (siANs; Figure 5A). Cultures were differentiated for 7 days in the presence of different trophic factors and pharmacological supplements selected to support neuronal differentiation. Glutamate-evoked calcium responses were then measured using Fluo-8 across a range of glutamate concentrations (**Figure 5A, B**). Response amplitude and concentration dependence varied with the supplements used during differentiation. LIF increased the maximal response relative to the factor-free control, and further addition of BDNF+NT-3 produced the largest maximal response, reaching 136 ± 8.1% of the LIF reference (**Figure 5C**). LIF+BDNF yielded a lower fitted EC₅₀ than LIF alone (0.182 and 0.698 μM, respectively). This assay provides proof of concept for high-throughput functional assessment in siAN cultures. Using maximal glutamate responsiveness as the criterion, LIF+BDNF+NT-3 was selected for further characterization.

**Figure 5.**
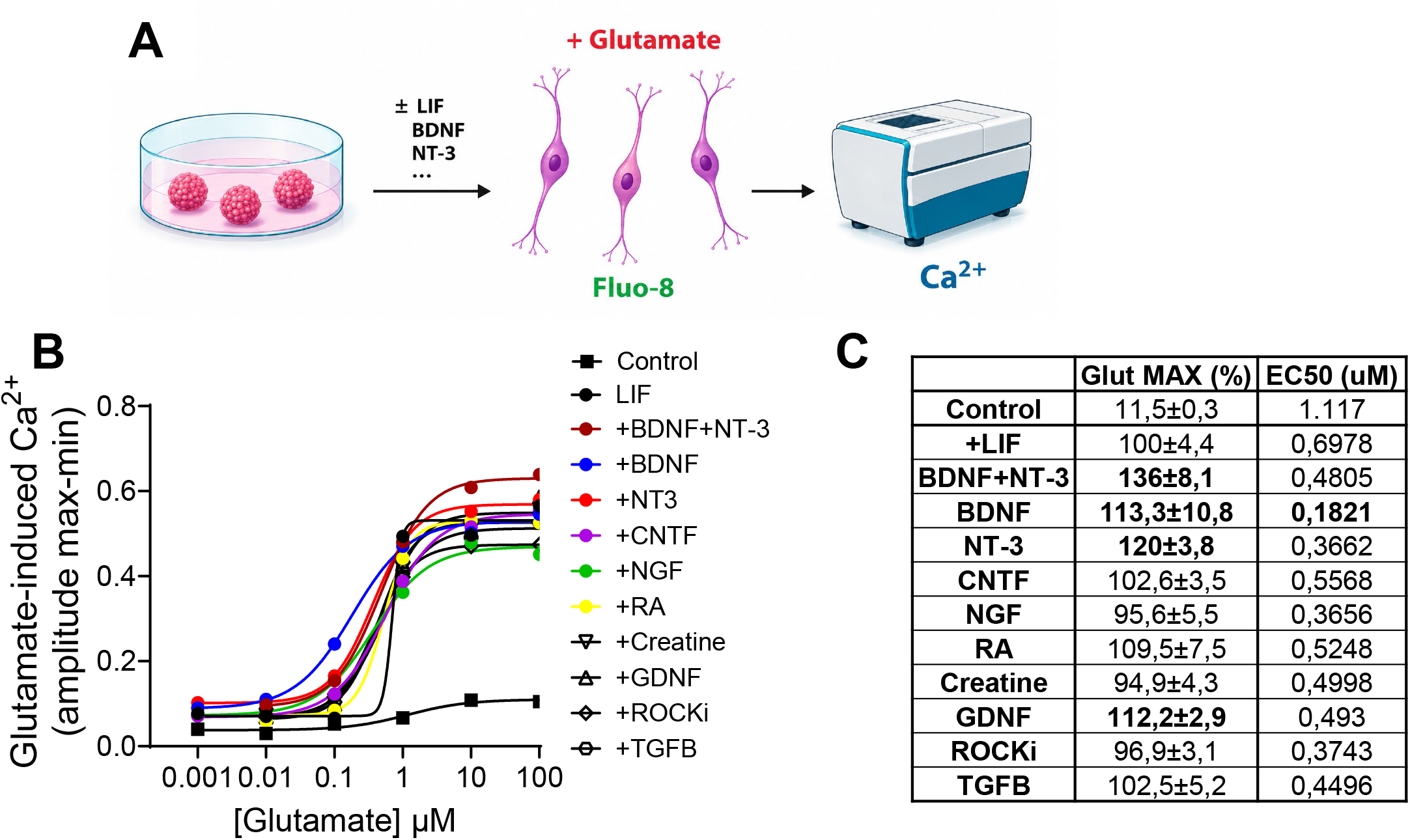
Functional screening of neurotrophic cues using glutamate-evoked Ca²⁺ responses in ANPG-derived neurons. **(A)** Schematic of the functional assay. Expanded ANPGs were differentiated in the presence of the indicated trophic factors or pathway modulators. After differentiation, cells were loaded with the Ca²⁺ indicator Fluo-8 and stimulated with glutamate to quantify evoked intracellular Ca²⁺ responses. **(B)** Glutamate concentration–response curves for ANPG-derived cells differentiated under the indicated conditions. Ca²⁺ responses were measured following stimulation with increasing concentrations of glutamate and fitted by nonlinear regression. **(C)** Pharmacological parameters derived from the concentration–response curves shown in (B), including maximal glutamate-evoked response (Glut MAX) and EC₅₀. Glut MAX values are expressed relative to the LIF condition, set to 100%. Calcium imaging was performed after 7 days of differentiation.

We further characterized neuronal differentiation under LIF+BDNF+NT-3 supplementation. Expanded progenitors expressed Nestin and SOX2, while cultures differentiated for 7 days contained βIII-tubulin-positive neuronal cells and S100-positive cells (**Supplementary Figure 3 A, C**). Transcriptomic comparison identified 987 genes enriched in expanded ANPGs and 1,570 enriched after differentiation (**Supplementary Figure 3B**). Expanded ANPGs showed enrichment of gene sets associated with translation and rRNA processing, whereas differentiation was accompanied by enrichment of neurogenesis, neuronal projections, axons and synapses (**Supplementary Figure 3D-F**).

### siANs support high-throughput screening for candidate neurogenic compounds

We next developed a siAN reporter assay for high-throughput screening of candidate neurogenic compounds. We generated a cryopreserved bank of more than 200 million expanded mouse ANPGs carrying a Tuba1-GFP reporter and used these cells to produce siAN cultures in 384-well plates (**Figure 6A**). The Tuba1a promoter was used to track neurogenesis [23]. This bank allowed cultures to be thawed and used within a defined passage range (passages 15–20), standardizing the source material for screening. GFP expression in differentiated cultures colocalized with βIII-tubulin, supporting its use for imaging neuronal cells and processes (**Supplementary Figure 4A, B**). LIF+BDNF+NT-3 increased GFP-positive cell yield to approximately 3.5-fold that of the control and provided a positive reference condition (**Figure 6B, C**).

**Figure 6.**
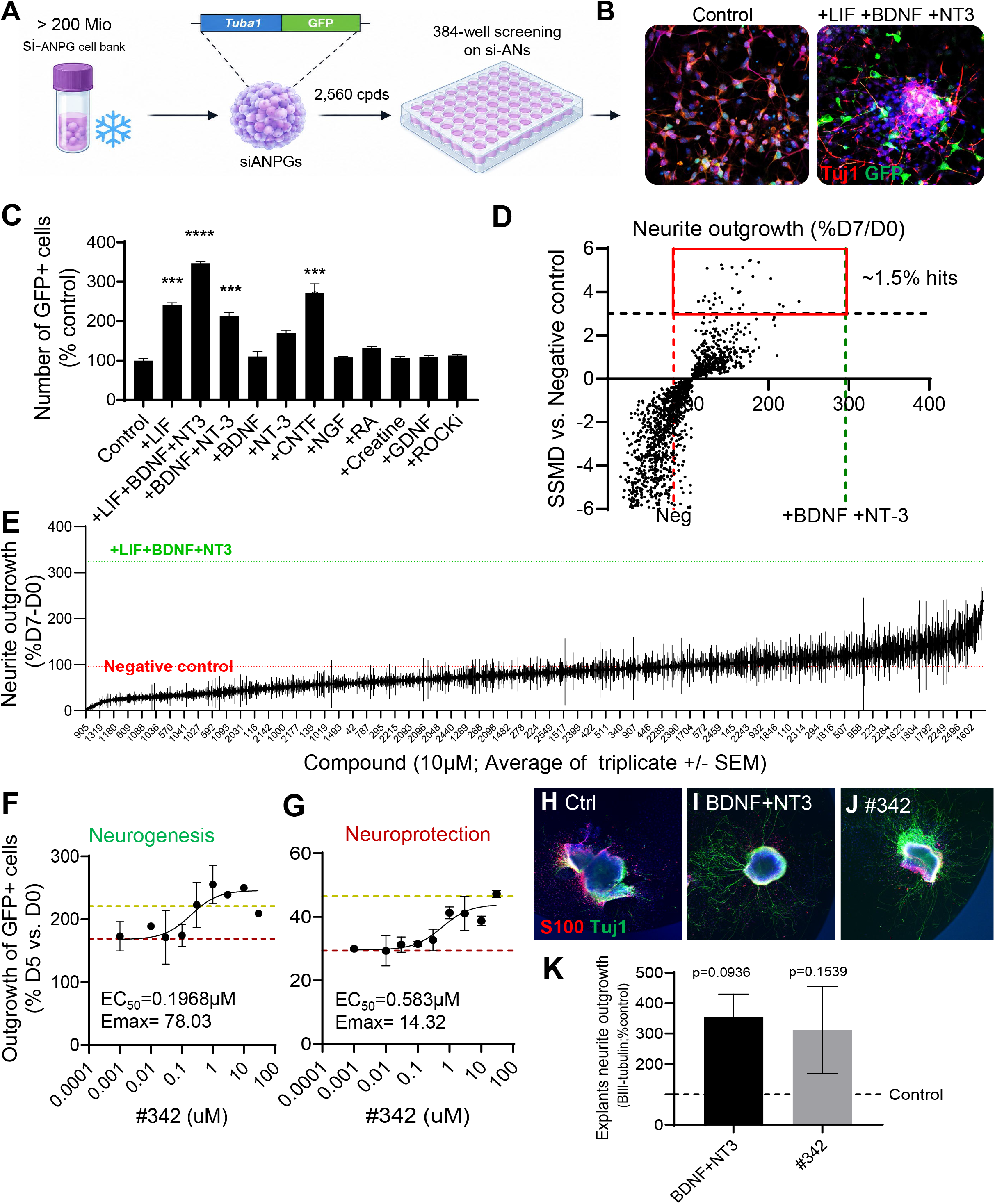
High-throughput Tuba1-GFP assay for auditory neurogenesis and hit validation. **(A)** Schematic of the screening workflow. Stemness-induced auditory neuroprogenitors (si-ANPGs) were transduced with a Tuba1-GFP reporter and expanded at large scale (>200 million cells) to generate a cryopreserved cell bank, enabling reproducible assays from thawed cells at matched passage. Cells were plated at 2,000 cells per well in poly-L-lysine/laminin-coated 384-well plates, with 10% proliferation medium during the first 24 hours. The medium was then replaced before compound addition, and GFP-positive neurite outgrowth was quantified on day 7, with LIF+BDNF+NT-3 serving as the positive control. **(B)** Representative confocal images of Tuba1-GFP-transduced si-ANs differentiated for 7 days in control medium or in the presence of LIF + BDNF + NT-3. GFP fluorescence reports neuronal differentiation, and immunostaining for βIII-tubulin (Tuj1) further confirms neuronal identity. Neurotrophin treatment increased both the number of GFP-positive cells and neurite outgrowth. **(C)** Quantification of GFP-positive cells after treatment with the indicated neurogenic factors and pathway modulators, expressed as percentage of control. Data are shown as mean ± SEM. Statistical significance is indicated in the graph. **(D)** Scatter plot summarizing primary screen results. Compounds were evaluated based on neurite outgrowth and SSMD relative to negative controls. The boxed region indicates candidate hits, corresponding to an overall hit rate of approximately 1.5%. **(E)** Rank-ordered distribution of screened compounds according to their effect on neurite outgrowth, expressed as %D7/D0. Negative and positive control benchmarks are indicated. **(F–G)** Dose-response validation of representative hit #342. **(F)** Neurogenesis assay showing increased neurite outgrowth of GFP-positive cells. **(G)** Cisplatin neuroprotection assay showing protection against cisplatin-induced neurite loss. EC₅₀ and Emax values are indicated in each panel. **(H–J)** Representative images from a secondary cochlear explant assay under control conditions (**H**), BDNF + NT-3 (**I**), or #342 (**J**). **(K)** Quantification of βIII-tubulin-positive neurite outgrowth in cochlear explants. Both BDNF + NT-3 and #342 showed a trend toward increased neurite outgrowth relative to control, but these effects did not reach statistical significance under the conditions tested.

Assay development included evaluation of seeding density, differentiation duration and an initial transition medium containing 10% proliferation medium (**Supplementary Figure 4C– F**). Although cultures differentiated without transition medium showed greater separation between positive and negative controls, poor survival of negative-control cells confounded this comparison. Addition of transition medium during the first 24 hours preserved control-cell survival. Under these conditions, 2,000 cells per well was the lowest seeding density yielding an SSMD above 3 at day 7 (SSMD 3.03) and was selected for screening.

The pilot screen was performed at 2,000 cells per well with a seven-day differentiation endpoint, allowing significant expression of neuronal markers in cultures **(Supplementary Figure 3)**. We screened 2,560 compounds at 10 μM in triplicate and quantified neurite outgrowth. Selection based on outgrowth and compound SSMD relative to the negative control identified approximately 1.5% of compounds as candidate hits (**Figure 6D, E**).

Compound 342 was selected for concentration-response follow-up and secondary assessment in primary tissue. The GFP-based outgrowth assay yielded a fitted EC_50_ of 0.197 μM, while an assay measuring protection against cisplatin-induced neurite loss yielded an EC_50_ of 0.583 μM (**Figure 6F, G**). We then assessed compound 342 in primary cochlear explants. Mean βIII-tubulin-positive neurite outgrowth was higher with compound 342 and BDNF+NT-3 than in control explants, although neither comparison reached statistical significance (p = 0.1539 and p = 0.0936, respectively; **Figure 6H-K**). This workflow establishes for the first time primary screening in renewable siAN cultures, concentration– response follow-up and secondary assessment in cochlear explants.

## DISCUSSION

Building on our established CHIR-DS protocol for scalable expansion of C57BL/6 ANPGs [14, 15, 24], this study characterizes the molecular and metabolic changes associated with induced stemness. The induced state combines reduced Id/Hes1 expression, increased accessibility at proneural bHLH motifs and progressive metabolic activation. Human fetal ANPG responsiveness provides proof of concept beyond mouse cultures. Neurons generated from expanded ANPGs (siANs) support glutamate-evoked calcium imaging and neurite-outgrowth screening. Together, these findings suggest potential molecular entry points for auditory neuronal regeneration and establish a renewable neuronal resource for functional studies and compound discovery.

### A transcriptional and chromatin response in ANPGs that involves the ID/bHLH circuit

bHLH transcription factors are key regulators of neural progenitor maintenance and lineage commitment, orchestrating the balance between quiescence, proliferation, and neuronal differentiation [25]. Among these, Ascl1 functions as a master regulator of neurogenesis, promoting cell cycle re-entry [17] and neural specification [26]. In quiescent ANPGs, omics profiling revealed a transcriptionally repressive landscape marked by elevated expression and accessibility of Hes1, Id1, Id2, Tle1, and Notch4, all known antagonists of bHLH activity that preserve quiescence and suppress neurogenic commitment [17] . Upon stemness induction using CHIR-DS (WNT activation with dual Smad inhibition) or CHIR+BMP (WNT activation with ALK2 inhibition), we observed increased ASCL1-associated chromatin accessibility, accompanied by enhanced neurosphere formation and ANPG proliferation. This derepression is likely mediated by the profound downregulation of Id family members (Id1, Id2), which are direct targets of Smad signaling and known inhibitors of bHLH function through sequestration of E-proteins [27]. ID downregulation would be expected to release E-protein partners such as E2A and HEB, enabling productive heterodimerization with proneural bHLH factors. Reduced Hes1 expression and accessibility, together with reduced accessibility at the corepressor locus Tle1, suggest an additional weakening of repression of proneural programs [17, 28, 29]. Notch inhibition has also been extensively investigated as a means of promoting supporting-cell proliferation and hair-cell regeneration in the mammalian cochlea [30]. In our sphere growth assay, further Notch inhibition with avagacestat or DAPT produced a modest increase in mean sphere growth, although this did not reach statistical significance (**Supplementary Figure 2**). Together, these observations suggest reduced ID/HES1-associated repression and increased accessibility at proneural bHLH motifs during ANPG stemness induction. Lower ID levels could increase the availability of E-protein partners, while changes at Hes1 and Tle1 suggest weaker transcriptional repression. This identifies the ID/bHLH network as a candidate link between BMP–SMAD inhibition and neurogenic competence.

### Metabolic adaptation during ANPG expansion

Stemness induction was accompanied by transcriptional enrichment of glucose metabolism, translation and growth-associated programs, followed by distinct metabolic responses during early and prolonged culture. Newly induced ANPGs showed modestly increased ECAR without increased OCR, consistent with an early enhancement of glycolytic activity. After prolonged expansion, both readouts were markedly elevated, indicating broader metabolic activation that included increased mitochondrial respiration. High basal respiration with limited additional respiratory reserve suggests sustained engagement of respiratory capacity, while increased acidification following mitochondrial inhibition indicates the ability to recruit compensatory glycolysis.

This profile is consistent with the energetic and biosynthetic demands of sustained ANPG expansion. Increased expression of translation-related genes could reflect greater demand for protein synthesis, while glucose metabolism could supply both energy and biosynthetic precursors. Enrichment of mTOR-and HIF1-associated gene sets further supports a growth-associated metabolic response. These observations align with coordinated cell-cycle and mitochondrial remodeling during neural stem-cell activation [31] and mirror well-established principles from other stem cell systems, where elevated glycolysis supports cellular expansion [32]. Notably, this shift coincided with transcriptional activation of canonical growth and nutrient-sensing pathways and components of the mTOR axis, further supporting a link between metabolic remodeling and the acquisition of stem-like features [33]. These findings are consistent with broader stem cell literature, where mTOR-HIF1α-MYC networks and glycolytic shifts support proliferation and self-renewal across neural and epithelial progenitors [31]. Together, these findings suggest that metabolic reprogramming is an integral component of the ANPG stemness transition.

### Cellular origin and human relevance of ANPG stemness induction

The cellular origin of ANPGs is central to understanding stemness induction and its regenerative implications. Several studies identify cochlear glia as a reservoir of latent neurogenic potential. Lang et al. (2015) showed that SOX2-positive glia in the adult auditory nerve possess neurosphere-forming and neural differentiation capacity, while neuronal injury stimulates proliferation of SOX2/SOX10-double-positive glia and reduces nuclear chromatin condensation [6]. Chen et al. (2021) further demonstrated that the SOX2-positive subset of cochlear glia has a high capacity for neuronal differentiation in culture and that small molecules promote acquisition of SGN-like features [9]. Evidence for a glial contribution to neuronal regeneration also extends to the adult cochlea in vivo. Using Sox10 lineage tracing, Wakizono et al. (2021) identified Schwann cells as a source of newly generated neurons following auditory nerve injury[34]. More recently, Wang et al. (2025) identified a FZD10-positive cochlear glial population whose proliferation and neuronal differentiation are enhanced by WNT signaling, further linking resident glial competence to a pathway targeted by our expansion protocol [11].

In this context, CHIR-DS may maintain or reactivate progenitor competence within glial-derived cells, with reduced ID/HES-associated repression and increased accessibility at proneural bHLH motifs providing candidate mechanisms for this response. Establishing this connection requires identification of the cells that initiate and sustain ANPG cultures. Lineage tracing initiated before culture, combined with single-cell profiling, would distinguish expansion of a pre-existing neurogenic subset from acquisition of neuronal competence by additional glial populations. Resolving the cellular origin would therefore link the molecular response characterized here to defined resident populations that could be targeted for endogenous auditory neuronal regeneration.

The human fetal experiments extend the relevance of the platform beyond mouse ANPGs. Human fetal ANPGs responded to the mouse-derived CHIR-DS protocol with increased sphere growth, continued propagation, maintenance of progenitor markers and subsequent expression of neuronal and glial markers. These results provide proof of concept for applying the expansion strategy to human auditory progenitors and support further evaluation of their regenerative potential.

### Platform application

Beyond regenerative studies, this work establishes expanded ANPGs as a renewable source for high-throughput neuronal assays. Building on the screening potential established for Phoenix cultures [24], the present 384-well, 2,560-compound campaign identified candidate enhancers of neurite outgrowth in differentiated siANs. A banked progenitor supply supports repeated assays and concentration–response testing feasible within a common culture system, addressing a major constraint of assays dependent on repeated primary tissue isolation. The approximately 1.5% candidate-hit fraction and follow-up of compound 342 demonstrate a route from compound screening to biological assessment. The reporter provides a scalable readout, while orthogonal measurements of neuronal identity, morphology and viability can distinguish effects on neurite extension, survival, differentiation or reporter activity. Repeated use of expanded cultures also offers a practical route to reduce fresh animal tissue requirements for primary screening.

Our cryopreserved bank of more than 200 million ANPGs enabled a standardized screening pipeline, illustrated here by a proof-of-principle screen of 2,560 compounds. This is almost six times the library previously examined in primary SGN cultures [35]. Triplicate measurements in standardized siAN cultures support primary hit selection, while the variability of SGN explants typically requires larger groups to distinguish treatment effects reliably. In our previous comparison of Phoenix-derived neurons and SGN explants, testing five neurotrophin doses involved 279 explants, with 22–94 explants per group, to detect significant differences in neurite outgrowth [24]. Published dissection protocols yield approximately 6–16 SGN explants per mouse when both cochleae are used [24, 36]. Applying the previously used range of 22–94 explants per group to 2,560 compounds would require approximately 3,500-40,000 mice, assuming 6–16 explants per mouse and excluding controls and preparation losses. Renewable ANPG cultures consequently offer a considerable 3R advantage by enabling broad screening while reserving adequately replicated explant experiments for prioritized candidates. siANs also support screening after one week of differentiation in 384-well plates, compared with more than ten weeks of preparation in a recent human inner ear organoid study [37]. Together with functional calcium imaging, these cultures support compound assessment across neuronal growth, protection and responsiveness, followed by evaluation of selected candidates in mouse explants or human organoids.

In our explant follow-up, mean neurite outgrowth was higher with compound 342 and the BDNF+NT-3 reference, although neither comparison reached statistical significance. We divided each explant into basal, middle and apical sections, whose differing responses to treatment may contribute to sample heterogeneity. The increase in mean outgrowth with compound 342 was comparable to that observed with the neurotrophin mix, supporting further assessment with a larger sample. More broadly, candidate evaluation could extend beyond outgrowth to hair-cell innervation and synapse formation. Expanded Lgr5-positive cochlear progenitors and optimized inner ear organoids provide complementary systems for generating sensory hair cells and studying ototoxicity[38, 39]. Vincent et al. demonstrated functional transmission at regenerated hair cell–SGN synapses in coculture, including refined release properties with older sensory epithelia[40]. Combining neuronal and sensory epithelial models would therefore offer a relevant next step for assessing whether compounds identified in siAN screens promote functional auditory neuronal repair.

## Conclusions

Molecular and metabolic characterization of the CHIR-DS-induced ANPG state identifies candidate molecular entry points for auditory neuronal regeneration. Genetic validation of the targeted pathways and responsiveness of human fetal ANPGs support the expansion strategy, while scalable siAN assays establish its practical value for neuronal studies and compound discovery.

## METHODS

### Study approval

All animal procedures were approved by the local veterinary office and the Commission for Animal Experimentation of the Canton of Geneva, Switzerland, authorization numbers GE66 and GE395. Procedures using human fetal material were performed with full approval by the Ethics Committee of the Geneva State, Switzerland, and following signed informed consent of the donors (authorization number 2024-00891).

### Spiral ganglion cell isolation and *in vitro* culture

Auditory neuroprogenitors were dissected from postnatal day 3-9 (P3-P9) C57BL/6J mice (Stock No: 000664) as previously described [14, 15]. Auditory neuroprogenitors (ANPGs) were isolated from the spiral ganglion, whereas otic progenitor cells (OPCs) were derived from the cochlear duct sensory epithelium following microdissection of the organ of Corti region. Both sexes were used without selection bias. Tissue dissociation was achieved following enzymatic digestion (StemPro™ Accutase™ Cell Dissociation Reagent; Thermo Fisher, Zug, Switzerland) for 10–15 min at 37 °C and mechanical pipetting trituration with a P1000 (10–15 up and downs). Following Accutase removal with a centrifugation step, the cell suspension was filtered with a 70-µm filter to eliminate undissociated tissue or bony parts. Isolated cells were maintained in culture with proliferation media consisting in DMEM:F12 with 15 mM HEPES buffer and 2 mM L-glutamine supplemented with 1× N2 and B27 supplement (Thermo Fisher, Zug, Switzerland), 1× penicillin streptomycin (100 U/mL) (Thermo Fisher, Zug, Switzerland), in the presence of basic fibroblast growth factor (bFGF; 10 ng/mL; ProSpec, Luzern, Switzerland), insulin like growth factor 1 (IGF1; 50 ng/mL; Cell Guidance Systems, Cambridge, UK), heparan sulfate (50 ng/mL; Sigma Aldrich, Buchs, Switzerland) and epidermal growth factor (EGF; 20 ng/mL; Cell Guidance Systems, Cambridge, UK), in ultra-low attachment six-well plates (Corning, Sigma Aldrich, Buchs, Switzerland). This growth factor combination is referred to throughout as GF. Following passaging or after initial dissociation, cells were manually counted using a FAST READ 102® (Milian, Plan-les-Ouates, Switzerland, reference 390497) hemocytometer, according to the manufacturer’s instructions. Cells were plated at 1 × 10^5^ cells/well and passaged twice a week using enzymatic digestion with Accutase™ followed by mechanical trituration as previously described.

### Pharmacological stemness induction

Four conditions were compared throughout the study. GF proliferation media (see paragraph above) was used as the reference condition. CHIR refers to GF supplemented with the WNT agonist CHIR99021 (3 µM; Axon Medchem, Groningen, The Netherlands). CHIR-DS refers to GF supplemented with CHIR99021 (3 µM) together with the TGFβ smad inhibitors LDN193189 (0.5 µM) and SB431542 (10 µM) (Axon Medchem, Groningen, The Netherlands), achieving dual smad inhibition. CHIR+BMP refers to GF supplemented with CHIR99021 (3 µM) and the ALK2 inhibitor K02288 (10 µM; Selleckchem, Munich, Germany). For the assessment of Notch inhibition, GF or CHIR media were supplemented with avagacestat or DAPT at 10 µM (Selleckchem, Munich, Germany). All compounds used in the study, with their respective targets, are listed in Supplementary Table 2. Compounds were prepared as 10mM except dorsomorphin and LY2109761 which concentration was 2mM stocks in DMSO, maintained at -80°C and diluted in media at the working concentration on the day of the experiment. The final DMSO concentration did not exceed 0.1% and was equivalent across all conditions of a given experiment.

### Multiplex miRNA mediated silencing of Alk2, Alk5 and Gsk3b

Synthetic miRNA minigenes were designed following our multiplex silencing technology as previously described [22], targeting *Alk2* (*Acvr1*), *Alk5* (*Tgfbr1*) and *Gsk3b*, corresponding to the BMP, TGFβ and GSK3β dependent regulation of the WNT pathway, respectively. Minigene sequences are available in Supplementary Table S3. Minigenes were cloned into pCWX-pGK-GFP and delivered to freshly isolated ANPGs by lentiviral transduction at MOI 10, maintained 5 weeks followed by selection with GFP using Biorad S3 cell sorter. Control cultures received control vector (pCWX-UBI-GFP). Knock-down efficiency was assessed by RT-qPCR. Neurosphere area was monitored over 30 days as described below, and cultures were followed along passages to assess long-term sphere formation.

### Automated analysis of ANPG neurosphere growth

ANPGs isolated from the P3–P9 old C57Bl6/J pup modiolus (see paragraph above) were plated at 10,000 cells/well in a 96 ultra-low attachment well plate, in 100 µL media per well. Wells containing ANPGs were organized in triplicates and placed in DMEM:F12, N2, B27 supplemented with growth factors (IGF1, EGF, bFGF and heparan sulfate; GF), with the WNT agonist CHIR99021 and/or the TGFβ smad inhibitors LDN193189 and SB431542, or with the additional pathway inhibitors listed in Supplementary Table 2. Phoenix ANPGs grown in GF conditions were used as positive control. Pictures of ANPG-derived neurospheres were performed twice a week using an ImageXpress plate reader over 38 days, and the size of the neurospheres was monitored using the MetaXpress software (v6.6.3.55, Molecular Devices, Biberach, Germany), with identical analysis parameters applied across all conditions. Experiments were repeated in at least three independent biological replicates.

### Human fetal ANPG isolation, culture and differentiation

Collection of human fetal inner ear spiral ganglion cells was done as previously described in [41]. Inner ears were isolated from aborted human fetuses ranging from week 10 to week 12 after conception. Dissociation and culture were performed as described above for mouse ANPGs. Human ANPGs were maintained under GF, CHIR (3 µM), DS (LDN193189 0.5 µM and SB431542 10 µM) or CHIR-DS, and neurosphere growth was monitored over 32 days using the acquisition and analysis settings described above. Cultures were enzymatically dissociated once a week and medium changed twice a week. For differentiation, mitogenic growth factors and reprogramming compounds were withdrawn and cells were plated on Matrigel 1/100 (hESC qualified; Corning, Buchs, Switzerland) coating for 7 days in differentiation media supplemented with LIF (10ng/ml), BDNF (25ng/ml) and NT3 (50ng/ml). Progenitor and neural markers were assessed by immunofluorescence and RT-qPCR, and glutamate-evoked calcium responses were recorded as described below. Three different donors were included in the study.

### Tuba1-GFP reporter ANPGs and cryopreserved cell bank

Expanded mouse ANPGs carrying a Tuba1a promoter driven GFP reporter were obtained following lentiviral transduction of stemness-induced ANPGs (pCWX-tuba1-GFP-PGK-bsd). Reporter expressing ANPGs were expanded in CHIR-DS conditions and frozen in DMEM:F12 with 15 mM HEPES buffer and 2 mM L-glutamine supplemented with 1× N2 and B27 supplement (Thermo Fisher, Zug, Switzerland) +10% DMSO at 4 million of cells per vial and stored in liquid nitrogen. The resulting bank consisted in more than 2 × 10^8^ expanded ANPGs. For all screening experiments, ANPGs were used between passage 15 and 20, unless stated otherwise in the figure legend. The relatively late passage usage allowed substantial material amplification and conservation through freezing, standardizing the source material across assay plates and screening runs. Colocalization of GFP with βIII-tubulin in differentiated cultures was verified by immunofluorescence.

### Differentiation of stemness-induced ANPGs into auditory neurons (siANs)

For differentiation, stemness-induced ANPGs were plated on laminin or Matrigel hESC qualified (**Supplementary Figure 2**) at ∼2–4 × 10^4^ cells/cm^2^ for 7 days with a medium change on day 3 or 4, following withdrawal of mitogenic growth factors and reprogramming compounds. Differentiation medium consists in DMEM:F12 with 15 mM HEPES buffer and 2 mM L-glutamine supplemented with 1× N2 and B27 supplement (Thermo Fisher, Zug, Switzerland), 1× penicillin streptomycin (100 U/mL) (Thermo Fisher, Zug, Switzerland), in the presence of LIF (Cell Guidance Systems, Cambridge, UK; 10 ng/mL), BDNF (Cell Guidance Systems, Cambridge, UK; 25 ng/mL) and NT-3 (ProSpec, Luzern, Switzerland; 50 ng/mL), or as stated otherwise in the figure legend. For the comparison of differentiation supplements (**Figure 5** and **Figure 6C**), the following factors were tested individually or in combination on a baseline of LIF (10 ng/mL): BDNF (50 ng/mL), NT-3 (50 ng/mL), CNTF (50 ng/mL), NGF (50 ng/mL), retinoic acid (10 µM), creatine (5 mM), GDNF (50 ng/mL), ROCK inhibitor (10 µM) and TGFβ (50 ng/mL) (CNTF, GDNF, ROCK inhibitor and TGFβ were purchased from Peprotech, NGF, retinoic acid and creatine from Sigma).

### Immunofluorescence and confocal microscopy

Stemness-induced ANPGs were differentiated for 7 days on Matrigel 1/100 (hESC qualified; Corning, Buchs, Switzerland) coated coverslips. For progenitors, undissociated neurospheres were left to attach on Matrigel 1/100 coated coverslips for 4 h in the incubator. Fixation, permeabilization and immunostaining of the samples were performed as previously described [15]. A list of antibodies used and dilutions is available in Supplementary Table S1. Nuclei were counterstained with DAPI. Confocal pictures were acquired on a LSM800 Airyscan microscope, and acquisition software ZEN Lite, and plate-based pictures were performed using an ImageXpress plate reader (Molecular Devices, Biberach, Germany). Acquisition settings were kept constant across conditions within an experiment, and pictures shown are representative of at least three independent experiments.

### RT-qPCR

RNA extraction was performed using a Qiagen RNA extraction minikit (Qiagen, Hombrechtikon, Switzerland, reference 74134), according to the manufacturer’s protocol. Reverse transcription was performed with Takara PrimeScript kit, with 200ng of RNA and quantitative PCR with SybrGreen on a QuantStudio12KFlex system from ThermoFisher Scientific. *TUBB3* and *GFAP* transcript levels were normalized to GAPDH, actin beta and EEF1 genes. Reactions were performed in 3 technical replicates on 3 independent biological replicates.

### Bulk RNA-seq sample preparation

Transcriptomic comparison was performed between ANPGs organoids, obtained from C57BL/6 treated with a combination of growth factors (GF; see above), GF+CHIR CHIR99021 3µM; GF+CHIR CHIR99021 3µM + LDN193189 0.5µM + SB431542 10µM; GF+CHIR CHIR99021 3µM +BMP (K02288) 10µM. Freshly isolated ANPGs were grown for 10 days in above mentioned conditions and RNA extraction was performed using a Qiagen RNA extraction minikit (Qiagen, Hombrechtikon, Switzerland, reference 74134), according to the manufacturer’s protocol. RNA concentration was determined using a Nanodrop spectrometer and RNA integrity was checked with the 2100 bioanalyzer (Agilent, Basel, Switzerland), following manufacturer’s instruction (RIN > 9). The RNA library was prepared as previously described and sequenced following Illumina TruSeq protocol [15]. For accurate quantitation of cDNA libraries, the fluorometric based system QuantiFluor™ dsDNA System was used (Promega, E2670). The size of final cDNA libraries was determined by using the dsDNA 905 Reagent Kit (Advanced Bioanalytical, DNF-905) exhibiting a sizing of 300 bp on average. Libraries were pooled and sequenced on the Illumina HiSeq 4000 sequencer (SE; 1 ×100 bp; 30–35 Mio reads/sample). The quality check was performed using FastQC [42].

### Bulk ATAC-seq sample preparation

Chromatin accessibility comparison was performed between ANPG organoids obtained from C57Bl/6 and treated with the same four conditions and for the same duration as described for RNA-seq. Freshly isolated ANPGs were grown for 10 days in the above-mentioned conditions, and 100,000 cells per sample were used for tagmentation. Tagmented DNA samples were prepared and purified using ATAC-seq Activ motif kit #53150. Libraries were pre-amplified for 5 cycles using indexed primers from Nextera kit (Illumina, FC-121-1031), and NEBNext High Fidelity PCR master mix (New England Biolabs, M0541S). Pre-amplified libraries served as input to determine the number of remaining cycles needed for the second round of PCR amplification to produce libraries. The remaining pre-amplified DNA was PCR amplified according to calculated Ct-value per sample and libraries were cleaned up by AMPure XP Beads (Beckman Coulter, A63880) at a 1.5× ratio. Library molarity and quality were assessed with Qubit (Thermofisher Scientific) and Tapestation using a DNA High sensitivity chip (Agilent Technologies, 5067-4626). Libraries were pooled at 2 nM and loaded for clustering on 1 lane of a Single-read Illumina Flow cell. Paired reads of 50 bases were generated using the TruSeq SBS chemistry on a NovaSeq 6000 sequencers.

### RNA-seq analysis

FastQ reads were mapped to the ENSEMBL reference genome (GRCm39.103) using STAR version 2.7.1a [43] with standard settings, except that any reads mapping to more than one location in the genome (ambiguous reads) were discarded (m = 1). A unique gene model was used to quantify reads per gene. Briefly, the model considers all annotated exons of all annotated protein coding isoforms of a gene creating a unique gene where the genomic regions of all exons are considered coming from the same RNA molecule and merged together. All reads overlapping exons and introns or only exons of each unique gene model were reported using featureCounts version 1.6.4 [44]. Counts of reads mapping only exons were transformed in RPKM and were used to filter low expressed genes (value lower than 1 RPKM). Raw counts of reads overlapping exons were used to call for DEGs. Library size normalizations and differential gene expression calculations were performed using the package edgeR [45] designed for the R software [46]. Only genes having a significant fold-change (Benjamini-Hochberg corrected p-value < 0.05) were considered for the rest of the RNA-seq analysis.

For heatmap and clustering, genes with abs (FC) ≥ 2 and *p*-value <0.05 (Fisher-exact test followed by Benjamini–Hochberg FDR correction) in the comparisons of interest were selected.

After hierarchical clustering of all selected DEGs using the function hclust in R, the number of clusters was assessed by cutting the dendrogram in 2 to 20 clusters using the function cutree in R. The final number of clusters was selected visually as the optimal number of clusters being the lowest number reporting correctly the variability of DEGs.

For gene ontology analysis, a GeneSetCollection R object including all Gene Ontology annotations obtained with biomaRt package for R was used [47]. A second GeneSetCollection R object was constructed by compiling all pathways and genesets from WikiPathways [48] KEGG [49] and Reactome [50] databases for both human and mouse. Gene ids not corresponding to the organism of the samples were translated to organism of interest homologous gene ids using biomaRt package for R. Enriched annotations within a set of genes were tested using the function GSEAGOHyperGParams of the Category [51] package designed for R. *P*-values were Benjamini-Hochberg corrected for multi-testing with the p.adjust function.

Enriched pathways in selected pairwise comparisons were used to generate enrichment plots (using enrichplot package [52] designed for the R software).

### ATAC-seq -Analysis

ATAC-seq libraries were analyzed following the validated ENCODE pipeline from Kundaje lab [53]. After adapter trimming with cutadapt version 1.9.1 [54] reads were mapped to the ENSEMBL reference genome (GRCm39.103) using bowtie2 version 2.2.6 [55]. Duplicated reads were marked with MarkDuplicates function from picard-tools version 1.126 [56] and later on removed using samtools view (-F 1804) version 1.2 [57]. Library complexities were assessed using bedtools version 2.26.0 [58] by calculating the ratios NRF (distinct/total), PBC1 (one read/distinct) and PBC2 (one read/two reads). Cross-correlations (NSC-Normalized Strand cross-correlation Coefficient and RSC-Relative Strand cross-correlation Coefficient) were calculated using the run_spp Rscript from Kundaje lab. For each biological replicate, two self-pseudoreplicates (containing half of the uniquely mapped reads) were generated following Kundaje lab recommendations. For each condition, pooled data of biological replicates and pooled-pseudoreplicates were created using zcat bash command. Reads on the forward and reverse strands were shifted by ^+^4 bp and 5 bp, respectively, to account for the 9 bp duplication created by DNA repair of the nick by Tn5 transposase. Broad and narrow peaks were called using macs2 version v2.1.2 [59] on each replicate, pseudoreplicate, pooled, and pooled-pseudoreplicate data, following Kundaje lab recommendations. Peaks overlapping genomic problematic regions [60] were removed using intersectBed from bedtools. Irreproducible discovery rate between each pair of biological replicates, pseudoreplicates, and pooled-pseudoreplicates was calculated using IDR 2.0.4 [61] Optimal final peaks were selected using 5% threshold for IDR. Finally, FRiP (fraction of reads in peaks) were calculated using intersectBed from bedtools.

For differentially accessible regions, for each condition, broad and narrow peaks were merged if they overlapped by at least 50% of the width of the most significant region using the package GenomicRanges [62] designed for R. Regions without overlap or overlapping by less than 50% were kept unchanged. Peaks were not merged between conditions assuming some redundancy in order to keep sensitivity as high as possible.

A list of all identified regions within all conditions was used to get read coverage in all biological replicates using featureCounts. Coverages were used as matrix input in edgeR for normalization and differential region accessibility calculations for all pairwise comparisons between conditions.

Localization of chromatin accessible regions (ChARs) was performed with the function findOverlaps from the package GenomicRanges designed for R, using the previously described unique gene model. ChARs overlapping promoters (−5000 bp/+5000 bp of each unique gene model TSS) by at least 1 bp were considered localized in promoter regions. The rest of the peaks which overlapped exonic regions by at least 1 bp were assigned to exons. Then, the rest of the peaks overlapping intronic regions by at least 1 bp were assigned to introns. Finally, the rest of the peaks were assigned to intergenic regions.

For heatmap and clustering, ChARs with abs(FC) ≥ 2 and *p*-value <0.05 (Fisher-exact test followed by Benjamini–Hochberg FDR correction) in the comparisons of interest were selected. Read coverage at each bp in selected ChARs in each biological replicate was assessed using coverageBed from bedtools. For each condition, ChAR coverages were normalized by the number of uniquely mapped reads in each replicate and the mean counts were reported and normalized in z-scores. After hierarchical clustering of all selected ChARs using the function hclust in R, the number of clusters was assessed by cutting the dendrogram in 2 to 20 clusters using the function cutree in R. The final number of clusters was selected visually as the optimal number of clusters being the lowest number reporting correctly the variability of ChARs.

To assign ATAC-seq peaks to genes, ChARs and genes were associated in pairs if they were separated by less than 500kb.

### Transcription factor binding site enrichment

Previously clustered ChAR sequences were extracted using the function subseq from the package Biostrings in R [63]. All identified ChARs not overlapping the clustered ChARs were used as background sequences. All sequences were resized by extension of 500bp upstream and downstream of their center. TF binding site enrichment vs. background sequences was tested for each cluster using findMotifs.pl script from HOMER (v4.9.1) [64]. Only TF binding site with a *p*-value below 10−4 (hypergeometric test) were represented. TCF4 in this analysis refers to the bHLH E-protein TCF4 (also known as E2-2), which is distinct from the WNT effector TCF7L2, sometimes called TCF4 in older literature.

### Aggregate ATAC-seq signal at genome-wide motif occurrences

In order to assess chromatin accessibility at bHLH binding sites independently of the accessibility at the *Tcf4* and *Ascl1* gene loci, genome-wide occurrences of the TCF4/E2-2 and ASCL1 motifs were identified in the GRCm39 reference genome using scanMotifGenomeWide.pl from HOMER v4.9.1, with the motif matrices reported in the cluster enrichment analysis. Normalized ATAC-seq coverage of each condition was then aggregated over a window of −0.5 to +0.5 kb around the center of every motif occurrence, using annotatePeaks. Each trace represents the mean signal across all genome-wide occurrences of a given motif for one condition. This measure is representative of the aggregate accessibility at bHLH binding sites throughout the genome and is not the same as the local accessibility at the *Tcf4* or *Ascl1* gene loci. Motif logos and HOMER motif enrichment q-values shown in Figure 3 correspond to the CHIR-DS condition relatively to control.

### Assessment of ANPG metabolism

Oxygen consumption rate (OCR) and extracellular acidification rate (ECAR) were measured in GF cultures, in newly induced CHIR-DS cultures at passage 1 and in long-term CHIR-DS cultures at passage 20, using a Seahorse XF Pro model; Agilent, Santa Clara, California, USA. Following Accutase dissociation, cells were seeded at 20000 cells/well in 96-well plate ultra-low attachment (Corning #3474) and left growing in suspension for 4 days before the assay. The fourth day, spheres were transferred to Seahorse XF Pro M cell culture microplate #103774-100, medium was replaced by Seahorse DMEM pH 7.4 medium #103575-100 supplemented with Glucose 17.5mM, Sodium Pyruvate 1mM and L-Glutamine 2mM (all from ThermoFisher Scientific). One hour before measurement, plates were equilibrated at 37 °C in a non-CO_2_ incubator. OCR and ECAR were recorded over measurement cycles of 3 minutes. For OCR measurements, Mito Stress kit #103015-100 kit was used. Following measurements were recorded: 3 for baseline, 4 after oligomycine addition (final concentration 2µM), 3 after FCCP addition (final concentration 2µM) and 3 after AntimycineA and Rotenone (final concentration 0.5µM). For ECAR measurements, Glyco Rate kit #103344-100 was used as following: 3 measurements for baseline, 4 measurements after AntimycineA and Rotenone (final concentration 0.5µM) and 6 after 2-deoxyglucose (final concentration 50mM) injection to confirm the glycolytic contribution to extracellular acidification. At the end of each assay, cells were lysed in situ and the total protein content per well was determined by BCA assay (Pierce BCA Protein Assay, Thermo Fischer Scientific). OCR and ECAR were normalized to the BCA protein measurement of the corresponding well. Each condition was assayed in 5 technical replicate wells over 3 independent biological replicates.

### Glutamate-evoked calcium imaging

Glutamate-evoked calcium responses were measured using a Fluo-8-based assay adapted from previously described Phoenix ANPG protocols [24]. Expanded stemness-induced ANPGs were dissociated with Accutase and plated at 30,000 cells per well in poly-ornithine/laminin-coated 96-well plates. Cells were differentiated for 7 days in mitogen-free medium supplemented with LIF, BDNF, and NT-3, or with neurogenic compounds as specified in the corresponding figure legend, with a medium change at day 3. At the end of differentiation, cells were loaded with Fluo-8 according to the manufacturer’s instructions and incubated for 45 min at 37°C. Calcium responses were acquired using an FDSS/µCELL Functional Drug Screening System. Cells were stimulated with increasing concentrations of glutamate ranging from 0.01 to 100 µM, using 1-log increments, and Ca²⁺-dependent fluorescence changes were recorded. Fluorescence kinetics were acquired for 15 min after the first glutamate addition, with one measurement every 0.5 s. Responses were quantified from background-subtracted fluorescence traces as the peak fluorescence amplitude relative to baseline. For dose-response analysis, response amplitudes were normalized, plotted against glutamate concentration, and fitted by nonlinear regression using a four-parameter logistic model to calculate EC₅₀ values.

### siAN neurite outgrowth assay in 384-well format

Tuba1-GFP ANPGs were thawed, used between passages 15 and 20, dissociated with Accutase™, and plated in poly-L-lysine/laminin-coated 384-well plates in 50 µL of differentiation medium per well, with 10% proliferation medium during the first 24 hours, as specified below. To establish assay conditions, GFP-positive neurite outgrowth was compared between control cultures and cultures supplemented with LIF (10 ng/mL), BDNF (50 ng/mL) and NT-3 (50 ng/mL). Seeding densities ranged from 2,000 to 10,000 cells per well without transition medium and from 1,000 to 5,000 cells per well with 10% proliferation medium during the first 24 hours. Readouts were acquired on days 3 and 7, using six replicate wells per condition. Pictures were performed using an ImageXpress plate reader and GFP-positive cells and neurite length were quantified using the MetaXpress software (v6.6.3.55, Molecular Devices, Biberach, Germany), applied identically across all wells and plates. Assay performance at each density and time point was assessed by the strictly standardized mean difference (SSMD), the Z′ factor and the signal-to-background ratio, calculated from the positive control (LIF+BDNF+NT-3) and negative control wells of each plate as SSMD = (µ_p_ − µ_n_) / √(σ_p_² + σ_n_²), Z′ = 1 − 3(σ_p_ + σ_n_) / |µ_p_ − µ_n_| and S/B = µ_p_ / µ_n_, where µ and σ are the mean and standard deviation of the positive (p) and negative (n) controls. A seeding density of 2,000 cells per well was selected as the lowest cell input yielding an SSMD above 3 at day 7 (SSMD 3.03), reducing the culture expansion time and incubator space required to generate material for screening. Addition of 10% proliferation medium during the first 24 hours preserved negative-control cell survival.

### Pilot screening of neurogenic compounds

The pilot screen was performed on siAN cultures seeded at 2,000 cells per well in poly-L-lysine/laminin-coated 384-well plates. Differentiation medium contained 10% proliferation medium during the first 24 hours and was then replaced with differentiation medium without proliferation medium before compound addition. Neurite outgrowth was assessed after 7 days of differentiation. A library of 2,560 compounds was screened at a final concentration of 10 µM in triplicates, with compounds added 24 hours after seeding and with a medium change right before compound addition as well as at day 4. Each plate contained negative control wells (vehicle, 0.1% DMSO) and positive control wells (LIF+BDNF+NT-3), organized in two different positions within the plate. Pictures were performed and neurite outgrowth quantified as described in the paragraph above. For each compound, neurite outgrowth at D7 was normalized to the neurite outgrowth of the same plate at D0 and a compound SSMD was calculated relatively to the negative control wells. Compounds were selected as candidate hits when SSMD > 3, corresponding to approximately 1.5% of the library.

### Concentration-response and cisplatin protection assays

Compound 342 was assessed in the GFP-based neurite outgrowth assay over a concentration range of 0.001, 0.003, 0.01, 0.03, 0.1, 0.3, 1, 3, 10, 30µM in 2 replicate wells, under the seeding and differentiation conditions used for the screen. For the cisplatin protection assay, siAN cultures were differentiated for 7 days and then exposed for 3 days to 10 µM cisplatin together with compound 342 tested over the same concentration range of 0.001–30 µM. Neurite length per well was quantified as described above and normalized to cisplatin-free and cisplatin-only control. Dose-response curves were fitted by nonlinear regression using a four-parameter logistic model to calculate EC_50_ values.

### Spiral ganglion explant culture and neurite outgrowth

Collection of mouse inner ear spiral ganglion cells for organotypic explant culture was done with P5–P7 C57Bl6/J pups, as previously described in [24]. Pups were euthanized by decapitation and the head surface was sectioned along the midsagittal plane. Brain and brainstem were removed, and both half-skulls were immersed in 10 mL of ice-cold Hanks balanced salt solution (HBSS) in a 100-mm Petri dish. The cochleae were removed under a stereo microscope (Nikon, Japan). The cartilaginous cochlear capsule was opened and the membranous labyrinth was exposed. The cochlear duct was separated from the modiolus (with the spiral ganglion cells) in one piece from base to apex using 5.5 forceps, by peeling it away. The organ of Corti and the stria vascularis appeared like two parallel ribbons adjacent to each other and were pulled apart to isolate the organ of Corti. Both spiral ganglion and organ of Corti were transferred separately into 1.5-mL Eppendorf tubes with ice-cold HBSS.

Spiral ganglion explants were divided in three parts (apical, medial and basal) with micro-scissors; organ of Corti explants were let entirely. Each explant was transferred in a well from a 96-well plate before medium addition. Wells were previously coated with Matrigel® (Corning #354277) diluted at 1/100 in medium and let 2 h at 37 °C; medium containing Matrigel was aspired just before adding explants, with no washing. Explants were cultured in DMEM:F12 with 15 mM HEPES buffer and 2 mM L-glutamine supplemented with 1× N2 and B27 supplement (Thermo Fisher, Zug, Switzerland) supplemented with compound 342 at 10µM, with BDNF and NT-3 at 25ng/ml as a positive reference, or with vehicle, as indicated in the figure legend. Half of the medium was changed after 3–4 days, and explants were fixed with PBS/paraformaldehyde 4% (ChemCruz #sc-281692) after one week in culture.

For immunostaining, explants were permeabilized with PBS/Triton (Sigma #X100) 0.2% and blocked with blocking buffer containing 1% BSA (Sigma #A3912) in PBS. For spiral ganglion explants, primary antibodies were S-100 (mouse, Sigma S2532) 1/500 and βIII-tubulin (rabbit, Biolegend #802001) 1/2000. For organ of Corti explants, primary antibodies were S-100 (mouse, Sigma S2532) 1/500 and myosin 7a (rabbit, Proteus 25-6790) 1/200. Secondary antibodies were donkey anti mouse-647; A31571 and donkey anti rabbit-555 (Thermofisher #A31572), all 1/1000. Explants were stained with DAPI (Applichem #A4099) 300 nM. Pictures were performed with an ImageXpress plate reader (Molecular Devices, Biberach, Germany). βIII-tubulin-positive neurite outgrowth was quantified using the MetaXpress software, with identical settings across all conditions. Explants were obtained from 3 independent litters, with 3 explants per litter (corresponding to basal, medial and apical cochlear turns) and final n=9.

### Statistical analysis

Data are presented as mean ± SEM unless otherwise specified. Statistical analysis was performed with GraphPad Prism software (v10.4.2). Depending on the experimental design, two-tailed t-tests, simple linear regression, one-way analysis of variance or two-way analysis of variance followed by appropriate post hoc tests were applied, as specified in the figure legends. Dunnett’s multiple-comparisons test was used for comparisons against a single control condition and Sidak’s multiple-comparisons test for selected pairwise comparisons. Dose-response data were fitted by nonlinear regression using a four-parameter logistic model and fitted EC_50_ values are reported. Values with p < 0.05 were considered as statistically significant. *p < 0.05, **p < 0.01, ***p < 0.001 and ****p < 0.0001. The number of independent biological replicates for each experiment is indicated in the corresponding figure legend. For statistical analysis of RNA-seq and ATAC-seq data, please refer to the corresponding paragraphs of the method section.

## Data availability

Bulk RNA-seq and ATAC-seq data have been deposited at GEO under the accession number GSE327831 and are publicly available as of the date of publication. All other data will be made available on reasonable request.

## Resource availability

Requests for further information and resources should be directed to and will be fulfilled by the lead contact, Francis Rousset. RNA-seq and ATACseq data have been deposited at GEO: GSE327831 and are publicly available as of the date of publication. All other data will be made available on reasonable request. Compounds screened in Figure 6 are reported using internal identifiers. Their chemical identities are withheld pending intellectual-property assessment.

## Acknowledgments

The authors would like to thank Mylène Docquier, from the Genomic platform of the University of Geneva for assistance with the transcriptomic data, Olivier Brun, from the bioimaging platform of the University of Geneva for assistance with the neurosphere growth assay and Sergei Startchik for assistance with tuba1-GFP+ neurite detection program.

## Author contributions

Conceptualization, F.R. and P.S.; investigation, F.R., S.S., M.S, L.P., S.L., S.I.; resources F.R. and P.S.; data curation F.R., S.L.; writing-original draft preparation, F.R.; writing-review and editing, F.R., P.S., M.S., L.O., D.D., S.L., S.B.; supervision, F.R., and P.S.; project administration, F.R., P.S.; funding acquisition, F.R., and P.S. All authors have read and agreed to the published version of the manuscript.

## Declaration of generative AI and AI-assisted technologies in the manuscript preparation process

During the preparation of this work the author(s) used AI in order to improve readability and language of this manuscript. The authors reviewed and edited the content as needed and take full responsibility for the content of the published article.

