## Supplementary material for "Deciphering auditory neuroprogenitor stemness for large-scale neuronal screening"

### Supplementary Figure 1

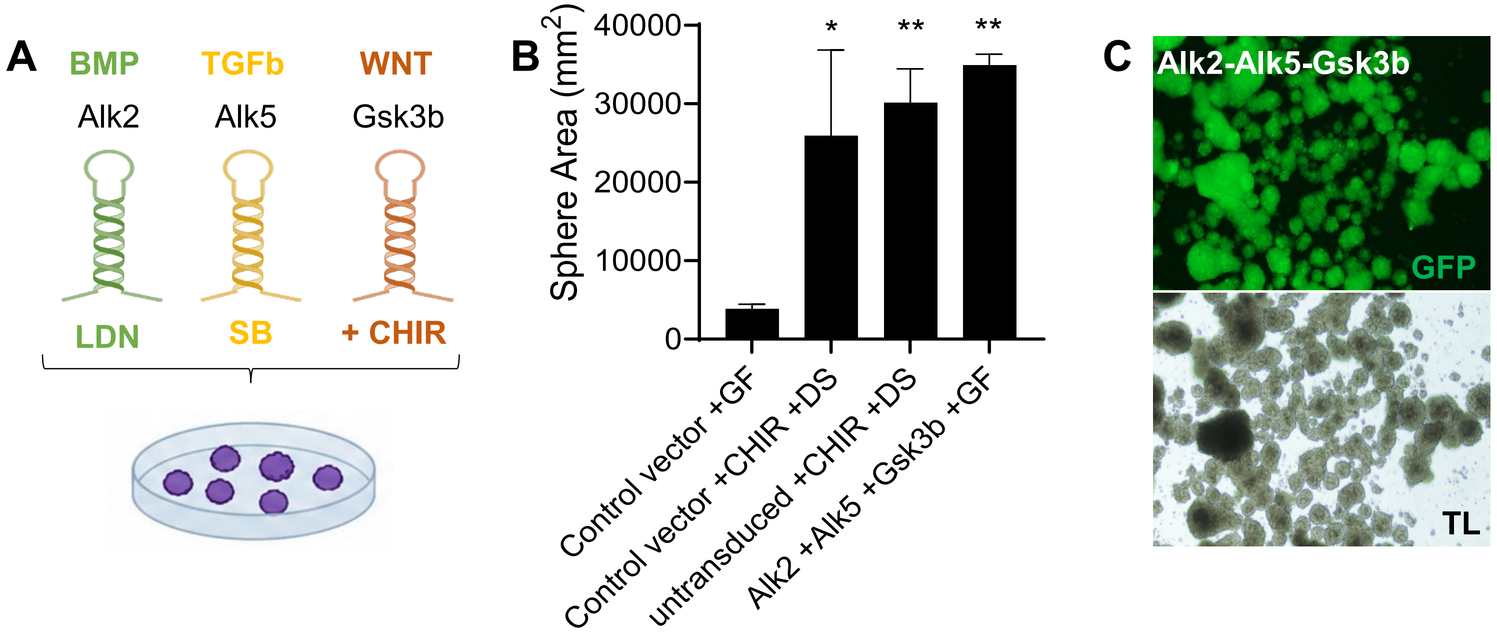


**Supplementary Figure 1. Mechanistic validation of CHIR-DS-induced stemness by multiplex genetic targeting of BMP, TGFβ, and WNT signaling**

**(A)** Schematic of the signaling nodes targeted to mechanistically validate the stemness-inducing effect of CHIR-DS observed in Figure 1. Pharmacological reprogramming combined WNT activation through GSK3β inhibition with CHIR99021, TGFβ pathway inhibition through ALK5 inhibition with SB431542, and BMP pathway inhibition through ALK2 inhibition with LDN193189. In parallel, the same signaling axes were targeted genetically using synthetic miRNA minigenes (TMGs) designed for multiplex gene silencing of *Alk2, Alk5,* and *Gsk3b*, based on the previously described TMG platform (Rousset et al., 2019).

**(B)** Quantification of sphere growth in low-stemness C57BL/6 ANPGs cultured for 30 days under the indicated conditions. Pharmacological treatment with CHIR + SB + LDN (CHIR-DS) significantly increased sphere area relative to growth-factor control. Control vector did not affect growth induction by CHIR +SB +LDN. Multiplex genetic silencing of *Alk2, Alk5,* and *Gsk3b* using the TMG approach similarly enhanced ANPG expansion. These results provide orthogonal mechanistic support for the contribution of BMP, TGFβ, and WNT signaling to the CHIR-DS-induced stemness phenotype. Data are presented as mean ± SEM; statistical significance is indicated in the graph.

**(C)** Representative images of ANPGs genetically reprogrammed with the *Alk2/Alk5/Gsk3b* TMG after long-term expansion. Upper panel, GFP fluorescence marking lentiviral transduction. Lower panel, corresponding transmitted-light image showing robust sphere formation and expansion.

Data are representative of three independent experiments.

### Supplementary Figure 2

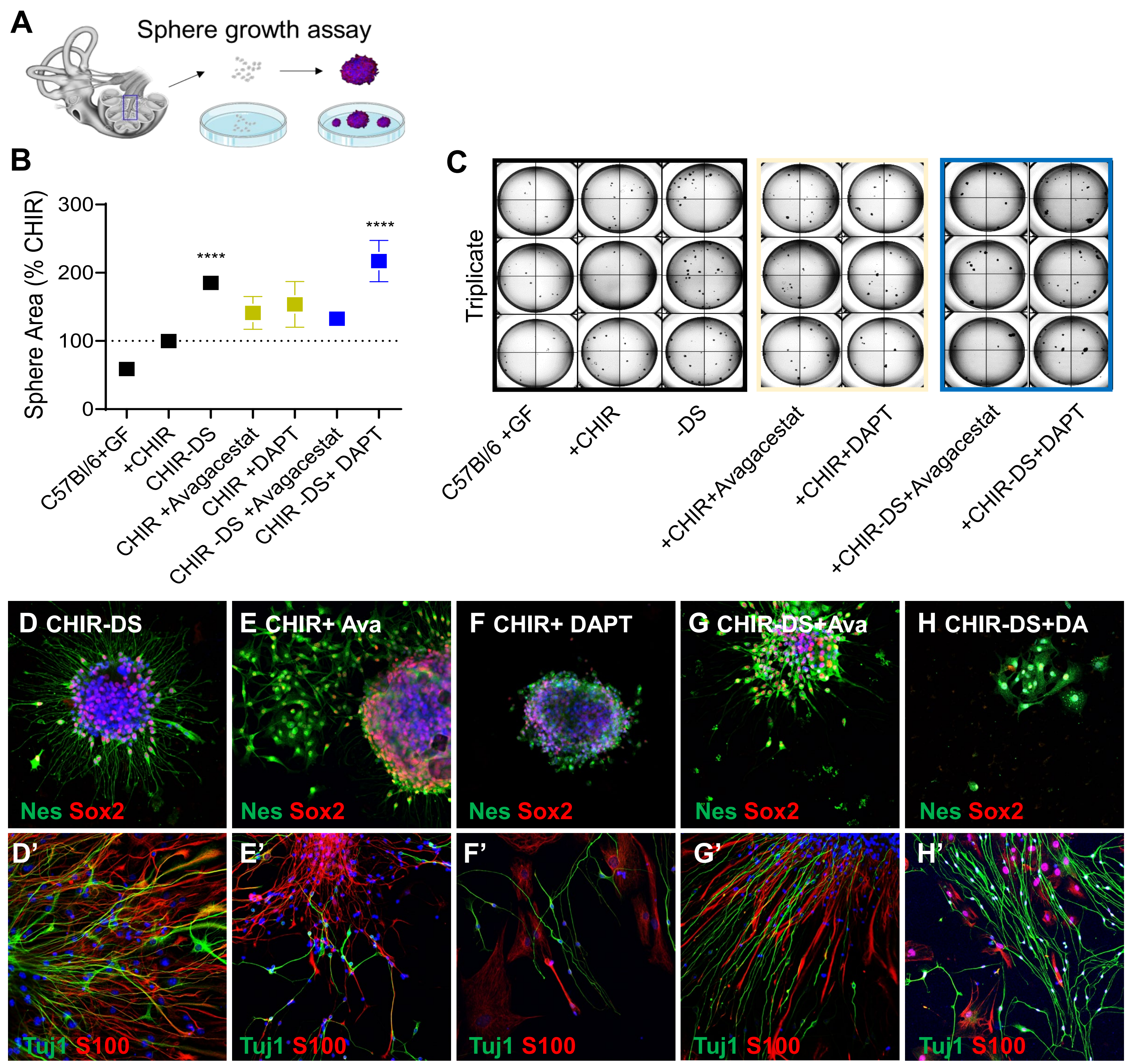


**Supplementary Figure 2. Impact of Notch inhibition on ANPG expansion and neural differentiation**

**(A)** Schematic of the ANPG sphere-growth assay used to assess the effects of Notch inhibition on stemness-associated expansion.

**(B)** Quantification of neurosphere area under the indicated conditions. Growth is expressed relative to the CHIR condition, set to 100%. Dual-SMAD inhibition (-DS) significantly increased sphere growth. Treatment with the γ-secretase inhibitors avagacestat or DAPT produced only modest changes in growth and did not significantly enhance expansion beyond the corresponding CHIR or CHIR-DS conditions. Data are presented as mean ± SEM; statistical significance is indicated in the graph.

**(C)** Representative macroscopic images of triplicate ANPG cultures under the indicated conditions, illustrating differences in neurosphere formation and growth.

**(D–H)** Representative immunofluorescence images of ANPG cultures treated with CHIR-DS (**D**), CHIR+avagacestat (**E**), CHIR+DAPT (**F**), CHIR-DS+avagacestat (**G**), or CHIR-DS+DAPT (**H**). Upper panels show expression of the neural progenitor markers Nestin (Nes, green) and SOX2 (red). Lower panels (**D′–H′**) show cultures after differentiation, stained for the neuronal marker βIII-tubulin (Tuj1, green) and the glial marker S100 (red), with nuclei counterstained with DAPI (blue).

### Supplementary Figure 3

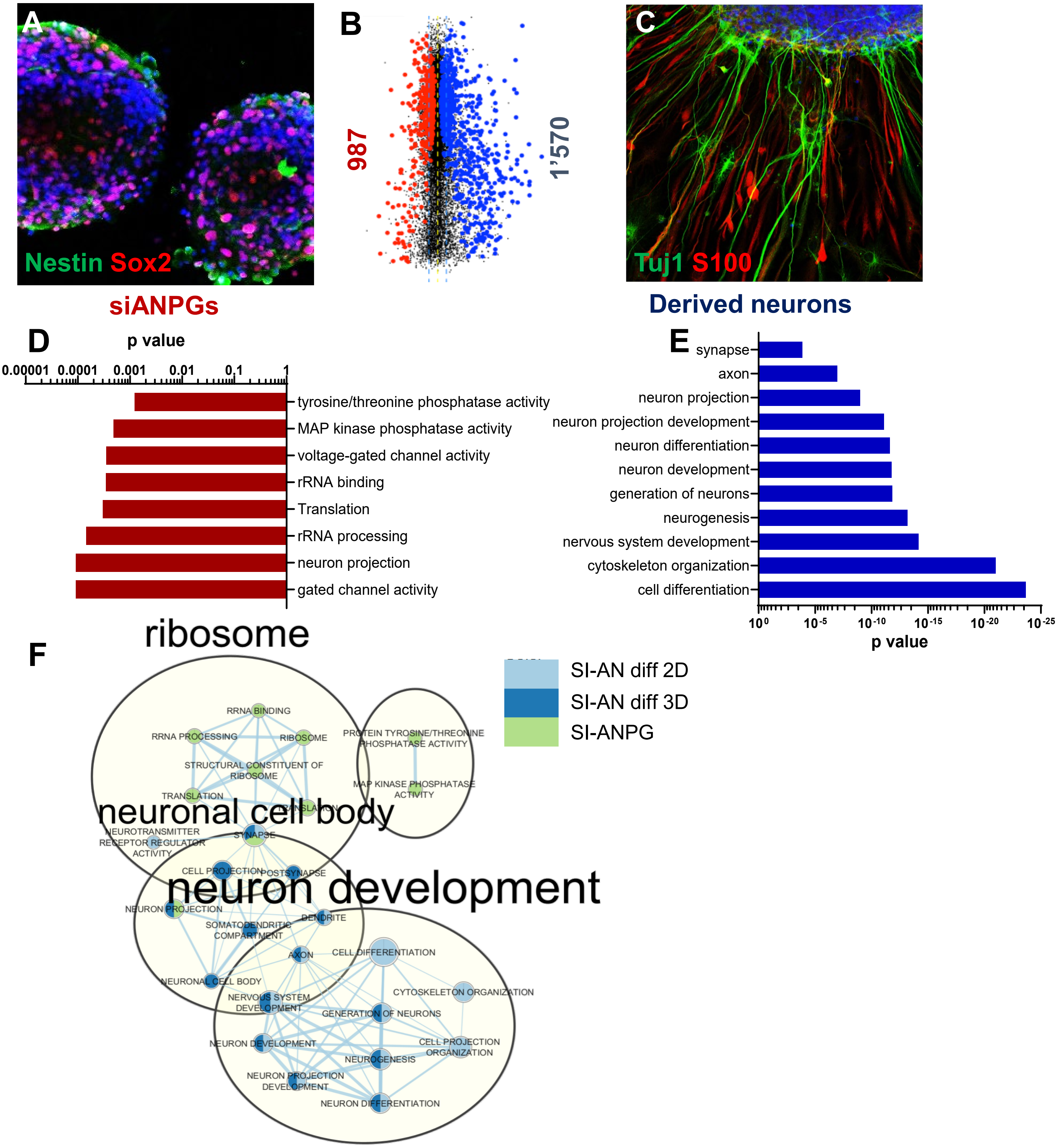


**Supplementary Figure 3. Transcriptional comparison of stemness-induced ANPGs and derived auditory neurons**
(**A**) Immunofluorescence images of expanded auditory neuron progenitor/glial cells (ANPGs) stained for Nestin (green) and Sox2 (purple).
(**B**) Dot plot showing the number and expression level of differentially expressed genes (DEGs) between expanded ANPGs and ANPG-derived auditory neurons after 7 days of differentiation in presence of LIF, NT-3 and BDNF.
(**C**) Immunofluorescence images of differentiated ANPG-derived neurons stained for the neuronal marker βIII-tubulin (Tuj1, green) and the glial marker S100β (red).
(**D, E**) Gene Ontology (GO) enrichment analysis highlighting biological processes significantly enriched in expanded ANPGs (**D**) and in ANPG-derived neurons (**E**).
(**F**) GO term network analysis illustrating shared and distinct biological processes between expanded ANPGs and differentiated ANPG-derived neurons. Nodes represent GO terms, and edges indicate shared genes between terms. Color coding denotes sample groups, including expanded ANPGs and ANPG-derived neurons differentiated in 2D and 3D conditions.
Data are representative of independent biological replicates. Statistical thresholds and enrichment parameters are described in the Methods section.

### Supplementary Figure 4

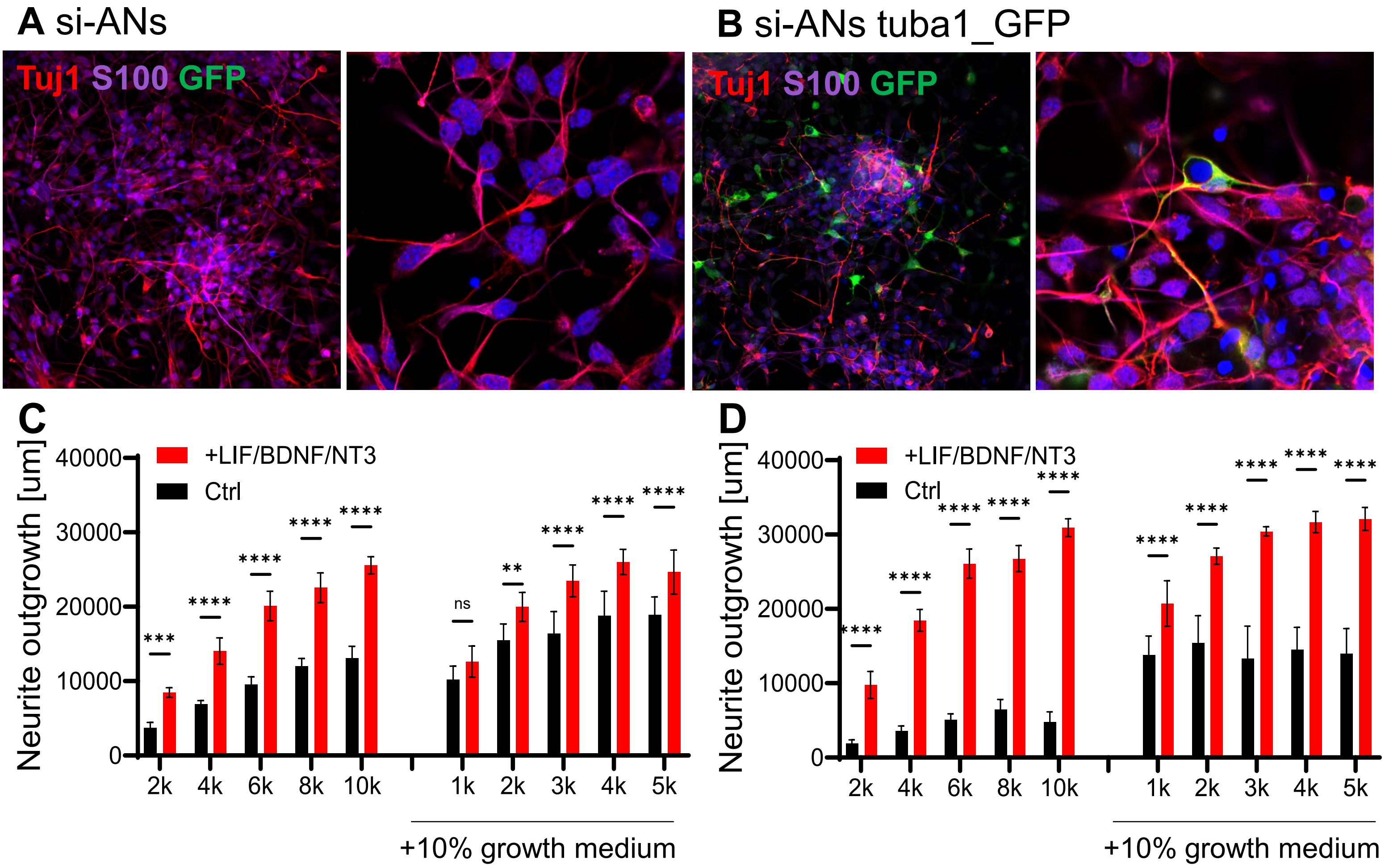


**E**

| D3 | Cells/well | Z′ | SSMD |
|---|---|---|---|
| | 2,000 | 0.126 | 4.85 |
| | 4,000 | 0.047 | 3.86 |
| | 6,000 | 0.128 | 4.66 |
| | 8,000 | 0.129 | 4.66 |
| | 10,000 | 0.334 | 6.29 |
| +Growth medium 10% | 1,000 | −3.843 | 0.87 |
| | 2,000 | −1.768 | 1.53 |
| | 3,000 | −1.167 | 1.93 |
| | 4,000 | −1.082 | 1.94 |
| | 5,000 | −1.806 | 1.50 |

**F**

| D7 | Cells/well | Z′ | SSMD |
|---|---|---|---|
| | 2,000 | 0.126 | 4.19 |
| | 4,000 | 0.577 | 9.31 |
| | 6,000 | 0.607 | 9.95 |
| | 8,000 | 0.543 | 9.21 |
| | 10,000 | 0.706 | 14.41 |
| +Growth medium 10% | 1,000 | −1.440 | 1.73 |
| | 2,000 | −0.232 | 3.03 |
| | 3,000 | 0.130 | 3.90 |
| | 4,000 | 0.227 | 5.17 |
| | 5,000 | 0.187 | 4.90 |

**Supplementary Figure 4. Validation of the Tuba1-GFP neuronal reporter and optimization of si-AN screening conditions**

**(A–B)** Representative immunofluorescence images of untransduced **(A)** and Tuba1-GFP-expressing **(B)** si-ANs after 7 days of differentiation in the presence of LIF, BDNF, and NT-3. Cells were stained for the neuronal marker βIII-tubulin (Tuj1, red), the glial marker S100B (magenta), and GFP (green). Nuclei were counterstained with DAPI (blue). GFP signal colocalizes with βIII-tubulin-positive neuronal cells and processes. Higher-magnification images (63×) are shown on the right of each panel.

**(C–D)** Quantification of GFP-positive neurite outgrowth after 3 **(C)** and 7 **(D)** days of differentiation at the indicated initial seeding densities. Black bars indicate basic differentiation medium, and red bars indicate supplementation with LIF, BDNF, and NT-3. Within each panel, cultures receiving 10% proliferation medium during the first day of differentiation (transition medium) are shown on the right. All cultures were plated in laminin/poly-L-lysine-coated wells. Bars represent mean ± SD from six replicate wells per condition.

**(E–F)** Assay performance at day 3 **(E)** and day 7 **(F)** for the corresponding seeding densities and medium conditions. Z′ factor and strictly standardized mean difference (SSMD) were calculated from GFP-positive neurite outgrowth in negative-control cultures maintained in basic differentiation medium and positive-control cultures supplemented with LIF, BDNF, and NT-3, using the means and sample SDs of six replicate wells per condition.

### Supplementary Table 1

| name | supplier and reference | dilution |
|---|---|---|
| ALEXA FLUOR 488 donkey anti rabbit | Life Technologies reference A21206 | 1/1000 |
| ALEXA FLUOR 555 donkey anti mouse | Life Technologies reference A31570 | 1/1000 |
| ALEXA FLUOR 555 donkey anti rabbit | Life Technologies reference A31572 | 1/1000 |
| ALEXAPLUS 488 donkey anti mouse | Life Technologies reference A32766 | 1/1000 |
| Ki67 | Abcam reference ab16667 | 1/100 |
| Myo7a | Proteus Bioscience reference 25-6790 | 1/200 |
| Nestin | Sigma reference ABD69 | 1/400 |
| Phalloïdin iFluor 488 | Abcam reference ab176753 | 1/1000 |
| S100 | Sigma reference S2532 | 1/500 |
| SOX2 | Santa Cruz reference sc365823 | 1/100 |
| Tuj1 | Biolegend reference 802001 | 1/2000 |

### Supplementary Table 2

| name | supplier and reference | stock solution | dilution | final concentration |
|---|---|---|---|---|
| K02288 | Selleckchem, reference S7359 | 10mM | 1/1000 | 10µM |
| Accutase | Invitrogen, reference A1110501 | | | |
| B27 supplement | Invitrogen, reference 17504044 | | 1/50 | |
| BDNF | CellGS, reference GFH1-100 | 10mg/ml | 1/400 | 25ng/ml |
| bFGF | Prospec, reference CYT-218 | 100mg/ml | 1/10000 | 10ng/ml |
| CHIR99021 | Axon kit, reference 5005 | 10mM | 1/3333 | 3µM |
| DMEM-F12 with HEPES and L-glutamine | Invitrogen, reference 31330038 | | | |
| EGF | Cell GS, reference GFH26-1000 | 100mg/ml | 1/5000 | 20ng/ml |
| Fibronectin | Millipore, reference FC010 | 1mg/ml | 1/200 | 0.005mg/ml |
| Heparane sulfate | AMSbio, reference AMS.GAG-HS01 | 100mg/ml | 1/2000 | 50ng/ml |
| IGF1 | Cell GS, reference GFH34-100, | 100mg/ml | 1/2000 | 50ng/ml |
| LDN193189 | Axon kit, reference 5005 | 10mM | 1/20000 | 0.5µM |
| LIF | Cell GS, reference GFH200-100 | 100mg/ml | 1/10000 | 10ng/ml |
| LY411575 | CellGS reference SM25 | 10mM | 1/1000 | 10µM |
| Matrigel hESC qualified | Corning, reference 354277 | | 1/100 | |
| N2 supplement | Invitrogen, reference 17502048 | | 1/100 | |
| NT3 | Prospec, reference CYT257 10UG | 10mg/ml | 1/200 | 50ng/ml |
| Penicilline/Streptomycine | Invitrogen, reference 15070063 | | 1/100 | |
| Poly-L-Ornithine | Sigma, reference P4957 | 0,01% | | 0,0015% |
| SB431542 | Axon kit, reference 5005 | 10mM | 1/1000 | 10µM |

### Supplementary Table 3

| TARGET | TMG sequence |
|---|---|
| Alk5 | GGTGATAGCAATGTCAGCAGTGCCTTTAAATCTCTGCCTCTCGGAACGTGAAGCCACAGATGGTTCCGAGAGACAGA<br>GATTTACAAGTAAGGTTGACCATACTCTAC |
| Alk2 | GGTGATAGCAATGTCAGCAGTGCCTTACAATTCCGTCTCCCTGAACCGTGAAGCCACAGATGGGTTCAGGGAAACGG<br>AATTGTCAAGTAAGGTTGACCATACTCTAC |
| Gsk3b | GGTGATAGCAATGTCAGCAGTGCCTTCTTTATCTCTGCTAACTTTCAGTGAAGCCACAGATGTGAAAGTTAGAAGAG<br>ATAAAGCAAGTAAGGTTGACCATACTCTAC |
